# Synergism of TNF-α and IFN-γ triggers inflammatory cell death, tissue damage, and mortality in SARS-CoV-2 infection and cytokine shock syndromes

**DOI:** 10.1101/2020.10.29.361048

**Authors:** Rajendra Karki, Bhesh Raj Sharma, Shraddha Tuladhar, Evan Peter Williams, Lillian Zalduondo, Parimal Samir, Min Zheng, Balamurugan Sundaram, Balaji Banoth, R. K. Subbarao Malireddi, Patrick Schreiner, Geoffrey Neale, Peter Vogel, Richard Webby, Colleen Beth Jonsson, Thirumala-Devi Kanneganti

## Abstract

The COVID-19 pandemic has caused significant morbidity and mortality. Currently, there is a critical shortage of proven treatment options and an urgent need to understand the pathogenesis of multi-organ failure and lung damage. Cytokine storm is associated with severe inflammation and organ damage during COVID-19. However, a detailed molecular pathway defining this cytokine storm is lacking, and gaining mechanistic understanding of how SARS-CoV-2 elicits a hyperactive inflammatory response is critical to develop effective therapeutics. Of the multiple inflammatory cytokines produced by innate immune cells during SARS-CoV-2 infection, we found that the combined production of TNF-α and IFN-γ specifically induced inflammatory cell death, PANoptosis, characterized by gasdermin-mediated pyroptosis, caspase-8-mediated apoptosis, and MLKL-mediated necroptosis. Deletion of pyroptosis, apoptosis, or necroptosis mediators individually was not sufficient to protect against cell death. However, cells deficient in both RIPK3 and caspase-8 or RIPK3 and FADD were resistant to this cell death. Mechanistically, the JAK/STAT1/IRF1 axis activated by TNF-α and IFN-γ co-treatment induced iNOS for the production of nitric oxide. Pharmacological and genetic deletion of this pathway inhibited pyroptosis, apoptosis, and necroptosis in macrophages. Moreover, inhibition of PANoptosis protected mice from TNF-α and IFN-γ-induced lethal cytokine shock that mirrors the pathological symptoms of COVID-19. In vivo neutralization of both TNF-α and IFN-γ in multiple disease models associated with cytokine storm showed that this treatment provided substantial protection against not only SARS-CoV-2 infection, but also sepsis, hemophagocytic lymphohistiocytosis, and cytokine shock models, demonstrating the broad physiological relevance of this mechanism. Collectively, our findings suggest that blocking the cytokine-mediated inflammatory cell death signaling pathway identified here may benefit patients with COVID-19 or other cytokine storm-driven syndromes by limiting inflammation and tissue damage. The findings also provide a molecular and mechanistic description for the term cytokine storm. Additionally, these results open new avenues for the treatment of other infectious and autoinflammatory diseases and cancers where TNF-α and IFN-γ synergism play key pathological roles.

## INTRODUCTION

The pandemic coronavirus disease 2019 (COVID-19), caused by severe acute respiratory syndrome coronavirus 2 (SARS-CoV-2), has caused over 1.2 million deaths in less than a year. SARS-CoV-2 infection results in a broad spectrum of clinical manifestations, including both asymptomatic cases and rapid fatalities (Petersen et al., 2020). The activation of the immune system and production of inflammatory cytokines are essential for the natural anti-viral immune responses (Crowe, 2017). However, hyperactivation of the immune system results in an acute increase in circulating levels of pro-inflammatory cytokines, leading to a “cytokine storm” (Tisoncik et al., 2012). While a mechanistic definition of cytokine storm is largely lacking, this storm is clinically characterized by systemic inflammation, hyperferritinemia, hemodynamic instability, and multi-organ failure.

During SARS-CoV-2 infection, patients present with systemic symptoms of varying severity which are associated with an aggressive inflammatory response and the release of a large amount of pro-inflammatory cytokines, or cytokine storm (Huang et al., 2020; Lucas et al., 2020; Yang et al., 2020). Studies have suggested a direct correlation between the cytokine storm and lung injury, multiple organ failure, and an unfavorable prognosis (Jose and Manuel, 2020; Mehta et al., 2020). Acute respiratory distress syndrome (ARDS) and systemic inflammatory response syndrome (SIRS) are other serious consequences of this cytokine storm (Ragab et al., 2020; Tisoncik et al., 2012). Understanding the underlying mechanisms driving the development of ARDS, SIRS, and multiple organ failure during cytokine storm is critical to develop targeted therapeutics against COVID-19 and other cytokine storm syndromes.

One possible mechanism linking cytokine storm to organ damage is the process of cell death. Among the programmed cell death pathways, pyroptosis, apoptosis, and necroptosis have been best characterized. Pyroptosis is executed by gasdermin family members through inflammasome activation–mediated caspase-1 cleavage of gasdermin D (GSDMD); caspase-11/4/5– or caspase-8–mediated cleavage of GSDMD; caspase-3–mediated cleavage of gasdermin E (GSDME); or granzyme A–mediated cleavage of gasdermin B (GSDMB) (He et al., 2015; Kayagaki et al., 2015; Orning et al., 2018; Sarhan et al., 2018; Shi et al., 2015; Wang et al., 2017; Zhou et al., 2020). Apoptosis is executed by caspase-3 and −7 following the activation of upstream initiator caspases caspase-8/10 or −9 (Elmore, 2007; Lakhani et al., 2006). Necroptosis is executed by RIPK3–mediated MLKL oligomerization (Dhuriya and Sharma, 2018; Giampietri et al., 2014; Murphy et al., 2013). Pyroptosis and necroptosis have been defined as inflammatory cell death processes, which are characterized as lytic forms of death that release cytokines and other cellular factors to drive inflammation and alert immune cells to a pathogenic or sterile insult, while apoptosis was historically considered immunologically silent. However, recent studies suggest that this is not always the case (Gurung et al., 2014; Gurung et al., 2016; Lukens et al., 2014; Malireddi et al., 2020; Malireddi et al., 2018). Depending upon the stimulus encountered, cells can experience extensive crosstalk among pyroptosis, apoptosis, and necroptosis called PANoptosis, which is inflammatory in nature (Christgen et al., 2020; Karki et al., 2020; Kesavardhana et al., 2020; Malireddi et al., 2020; Malireddi et al., 2018; Malireddi et al., 2019; Samir et al., 2020; Zheng et al., 2020a; Zheng et al., 2020b).

Currently, there is a critical shortage of proven treatment options for the ongoing COVID-19 pandemic, leading to an urgent need to understand the pathogenesis of multi-organ failure and lung damage in these patients. Given that cytokine storm is associated with severe inflammation and damage of vital organs during COVID-19, and that a detailed molecular pathway defining cytokine storm is lacking, gaining mechanistic understanding is critical to develop therapeutics. To date, there is no concrete evidence whether any of the cytokines induced in COVID-19 cause inflammatory cell death, and the cellular and molecular mechanisms employed by these cytokines in driving the pathophysiology of this pandemic disease remain unclear.

In this study, we evaluated the role of pro-inflammatory cytokines that are highly upregulated in patients with COVID-19 in inducing inflammatory cell death, inflammation, tissue and organ damage, and mortality. Our study shows that the specific combination of TNF-α and IFN-γ is critical for these processes. We found that inhibiting TNF-α and IFN-γ protected against death in SARS-CoV-2 infection and models of sepsis, hemophagocytic lymphohistiocytosis (HLH), and cytokine shock, suggesting that this pathway can be applicable beyond COVID-19 in infectious and inflammatory diseases where TNF-α and IFN-γ–mediated inflammatory cell death drive the pathology.

## RESULTS

### Inflammatory cytokines are elevated in patients with COVID-19, and synergy between TNF-α and IFN-γ specifically induces cell death

Studies of patients with COVID-19 have reported associations between disease severity and an influx of innate immune cells and inflammatory cytokines (Hadjadj et al., 2020; Huang et al., 2020; Lucas et al., 2020). To determine the pro-inflammatory cytokines that are the most highly upregulated during SARS-CoV-2 infection, we re-analyzed a publicly available dataset for circulating cytokines from healthy volunteers and patients with moderate or severe COVID-19 (Lucas et al., 2020). We observed that levels of circulating IL-6, IL-18, IFN-γ, IL-15, TNF, IL-1α, IL-1β, and IL-2 were the most upregulated in patients with moderate or severe COVID-19 **(Figure 1A)**. This is consistent with our previous results with MHV, a murine coronavirus (Zheng et al., 2020b), and several other studies in patients with COVID-19 (Blanco-Melo et al., 2020; Del Valle et al., 2020; Huang et al., 2020). Additionally, SARS-CoV-2–infected peripheral blood mononuclear cells (PBMCs) obtained from healthy donors showed increased production of pro-inflammatory cytokines **(Figure 1B)**.

**Figure 1.**
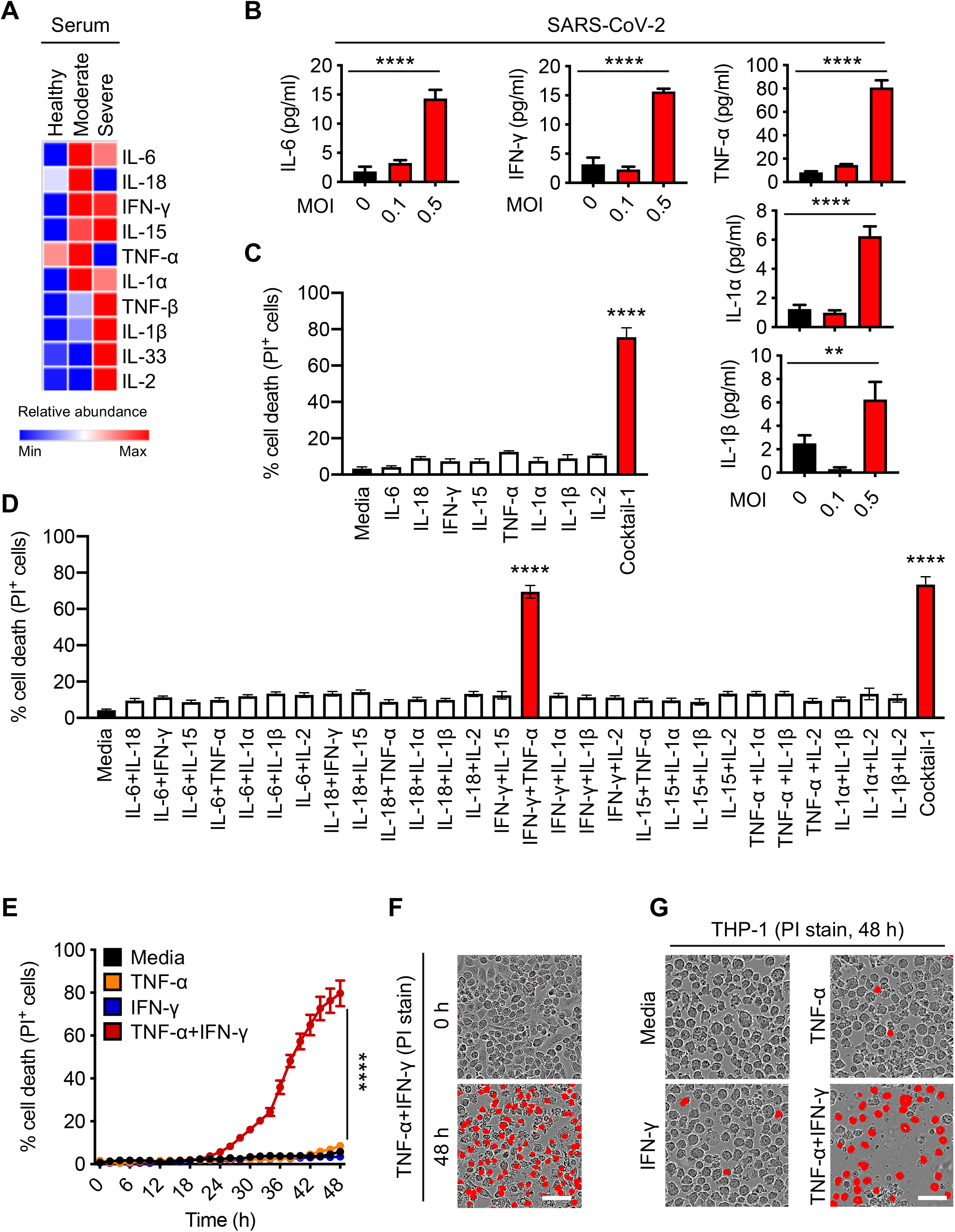
Pro-inflammatory cytokines are increased in COVID-19, and co-treatment of TNF-α and IFN-γ induces cell death. (A) Heatmap depicting the levels of pro-inflammatory cytokines in serum of patients with COVID-19 and healthy people (Lucas et al., 2020). (B) Pro-inflammatory cytokines released from PBMCs infected with SARS-CoV-2. (C) Percent of BMDMs that are dead 48 h after cytokine treatment using the IncuCyte imaging system and propidium iodide (PI) staining. “Cocktail-1” contained all 8 cytokines (IL-6, IL-18, IFN-γ, IL-15, TNF-α, IL-1α, IL-1β, and IL-2). (D) Percent of BMDMs that are dead 48 h after treatment with the indicated combination of cytokines. (E) Real-time analysis of cell death in BMDMs treated with the indicated cytokines. (F, G) Representative images of cell death in BMDMs (F) and THP-1 cells (G) after 48 h of the indicated treatments. Scale bar, 50 μm. Data are representative of at least three independent experiments. \*\**P* < 0.01; \*\*\*\**P* < 0.0001. Analysis was performed using the one-way ANOVA (B–D) or the two-way ANOVA (E). Significance asterisks in C and D indicate the comparison to the media-treated control. Data are shown as mean ± SEM (B–E). See also Figure S1.

A direct correlation between systemic increases in pro-inflammatory cytokines and lung injury, multiple organ failure, and unfavorable prognosis in patients with severe COVID-19 has been suggested, and uncontrolled cytokine signaling can contribute to multi-organ damage (Jose and Manuel, 2020; Mehta et al., 2020; Ragab et al., 2020). To understand the ability of the pro-inflammatory cytokines released during SARS-CoV-2 infection to induce cell death, we treated bone marrow-derived macrophages (BMDMs) with the cytokines that were the most highly upregulated in the circulation of patients with COVID-19 and in PBMCs infected with SARS-CoV-2. None of the cytokines individually induced high levels of cell death in BMDMs at the concentration used **(Figure 1C)**. However, the combination of all these cytokines (IL-6, IL-18, IFN-γ, IL-15, TNF-α, IL-1α, IL-1β, and IL-2; Cocktail-1) robustly induced cell death, suggesting that synergistic cytokine signaling is required for this process **(Figure 1C)**. To determine the specific cytokines involved in this synergy, we prepared all possible combinations of two cytokines. Out of the 28 combinations tested, only the combination of TNF-α and IFN-γ induced cell death to a similar extent as the cytokine cocktail (Cocktail-1) did **(Figure 1D)**. Treating with Cocktail-2, which lacked TNF-α and IFN-γ, failed to induce similar levels of cell death **(Figure S1A)**. TNF-α or IFN-γ might synergize with Cocktail-2 to induce cell death. However, addition of either TNF-α or IFN-γ individually to Cocktail-2 still failed to induce cell death, further supporting that synergism between TNF-α and IFN-γ is critical in inducing cell death **(Figure S1A)**. We then evaluated whether IFN-γ could be replaced with other IFNs that engage type I or type III IFN pathways, which are also important for immune regulation. Combining TNF-α with IFN-α, IFN-β, or IFN-λ failed to induce high levels of cell death **(Figure S1B)**, suggesting that type II IFN signaling specifically is crucial to co-ordinate with TNF-α signaling to induce cell death. The dynamics of cell death could be affected by the concentration of cytokines chosen. To confirm that the observed cell death was not an artifact of the cytokine concentration, we analyzed the effect of changing the concentrations of the cytokines in Cocktail-2 and found that increasing the concentration up to 10-fold still failed to induce cell death **(Figure S1C)**. Conversely, the kinetics of cell death induced by TNF-α and IFN-γ co-stimulation were proportional to the concentrations of TNF-α and IFN-γ **(Figure S1D)**. Similar to the cell death we observed in murine BMDMs **(Figures 1E and 1F)**, we also found that TNF-α and IFN-γ treatment induced robust cell death in the human monocytic cell line THP-1 and primary human umbilical vein endothelial cells (HUVEC) **(Figures 1G and S1E)**.

To understand the timing of TNF-α and IFN-γ release during the disease course of COVID-19, we analyzed another public dataset (Silvin et al., 2020) and found that increased production of TNF-α peaked in patients with moderate COVID-19, while production of IFN-γ was increased in patients with moderate disease and peaked in patients with severe COVID-19 (**Figure S1F)**. Additionally, the increased production of TNF-α and IFN-γ observed in patients with COVID-19 can be contributed by multiple cell types. To determine the contribution of immune cells in the production of TNF-α and IFN-γ, we re-analyzed publicly available single-cell RNA-seq data from PBMCs obtained from healthy donors and patients with mild or severe COVID-19 (Lee et al., 2020b). While macrophages from the PBMCs of patients with COVID-19 showed increased expression of TNF-α and IL-1α, NK cells and CD8^+^ T cells had increased expression of IFN-γ compared with healthy donors **(Figure S1G)**.

### TNF-α and IFN-γ shock *in vivo* mirrors COVID-19 symptoms

Patients with COVID-19 who require ICU supportive care often present with ARDS, acute cardiac injury, and acute kidney injury (Fan et al., 2020). These symptoms can be fatal and have been associated with cytokine storm. To examine whether TNF-α and IFN-γ can induce COVID-19– related symptoms, we developed a murine model of TNF-α and IFN-γ shock. Similar to our *in vitro* data, where treatment with TNF-α or IFN-γ alone failed to induce cell death, administration of TNF-α or IFN-γ individually did not cause significant mortality in mice. However, treating with the combination of TNF-α and IFN-γ led to synergistic mortality **(Figure 2A)**, indicating that the TNF-α and IFN-γ–mediated cell death may be associated with mortality. We then examined which cell types and organs were affected by the TNF-α and IFN-γ shock. We observed an increased influx of inflammatory cells in the lamina propria of the intestine of mice treated with TNF-α and IFN-γ compared with PBS-treated mice **(Figures 2B and S2A)**. Similarly, lungs from TNF-α and IFN-γ–treated mice showed septal thickening due to the accumulation of neutrophils in capillaries **(Figure S2B)**. Also, the incidence of caspase-3– and TUNEL–positive intestinal crypts and caspase-3–positive lung cells was increased in TNF-α and IFN-γ–treated mice compared with PBS-treated mice **(Figures 2B, S2B–S2D)**, suggesting that TNF-α and IFN-γ induce lung and intestinal damage and cell death. The increased cell death in the TNF-α and IFN-γ–treated mice was further confirmed by the presence of elevated serum LDH levels **(Figure 2C)**.

**Figure 2.**
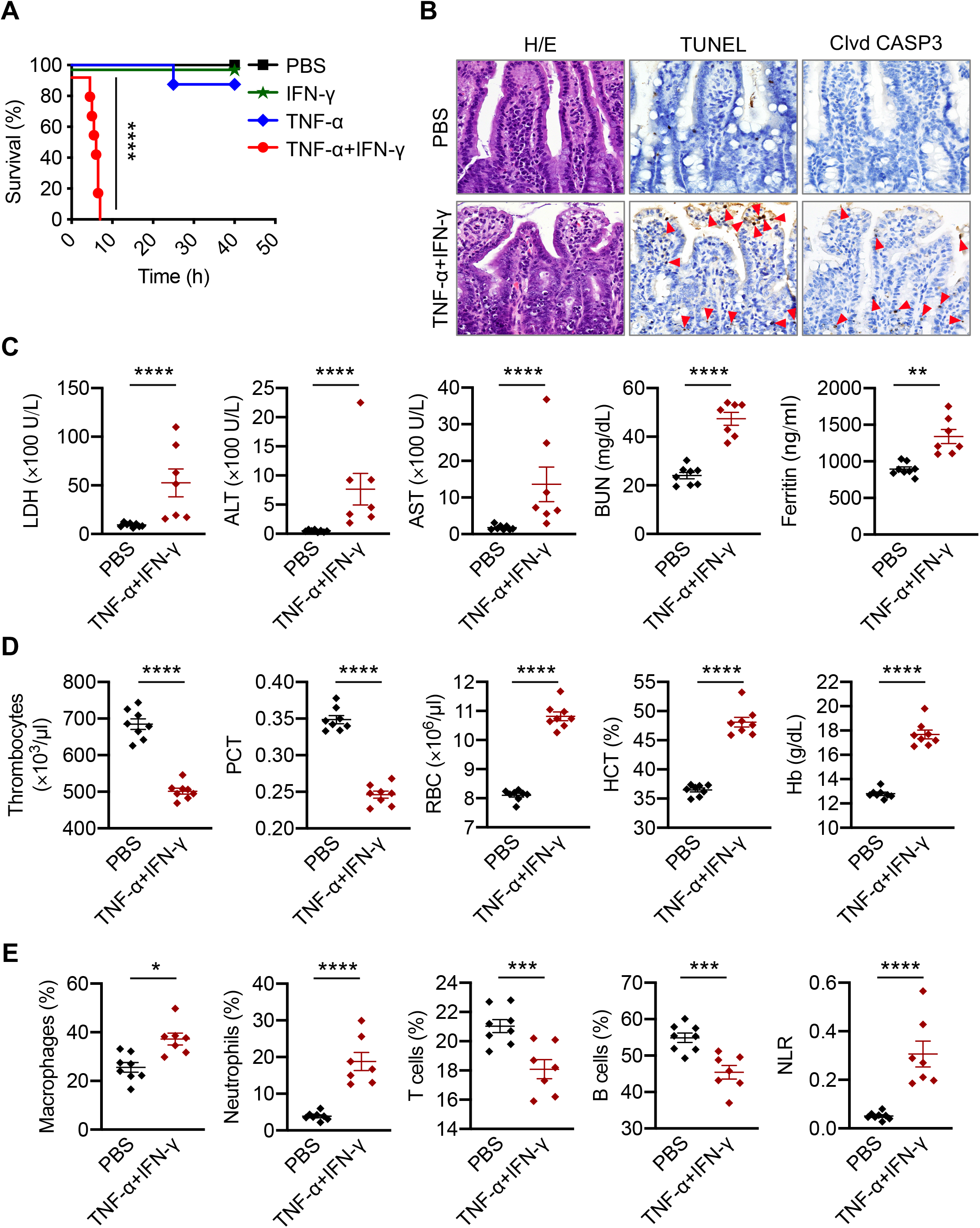
Cytokine shock by TNF-α and IFN-γ mirrors COVID-19 symptoms. (A) Survival of 6-to 8-week-old WT mice after i.p. injection of PBS (n = 10), IFN-γ (n = 12), TNF-α (n = 15), or TNF-α+IFN-γ (n = 15). (B) H/E staining, TUNEL, and cleaved caspase-3 (Clvd CASP3) immuno-staining of colon samples from PBS or TNF-α+IFN-γ injected mice for 5 h. Red arrows indicate stained cells. (C–E) Analysis of (C) serum levels of LDH, ALT, AST, blood urea nitrogen (BUN), and ferritin; (D) the number of thrombocytes, plateletcrit (PCT), RBC count, hematocrit (HCT), and hemoglobin (Hb) concentration in the blood; and (E) the percentage of macrophages, neutrophils, T cells, and B cells and the neutrophil-to-lymphocyte ratio (NLR) in the blood of PBS or TNF-α and IFN-γ injected mice for 5 h. Data are representative of at least three independent experiments. \**P* < 0.05; \*\**P* < 0.01; \*\*\**P* < 0.001; \*\*\*\**P* < 0.0001. Analysis was performed using the survival curve comparison (log-Rank [Mantel-Cox] test) (A) or the *t* test (C– E). Data are shown as mean ± SEM (C–E). See also Figure S2.

We next sought to determine whether TNF-α and IFN-γ shock could mimic the laboratory abnormalities observed in patients with COVID-19. In patients who succumbed to COVID-19, the levels of ALT, AST, blood urea nitrogen (BUN), and ferritin have been shown to increase until death (Ghahramani et al., 2020; Huang et al., 2020). Similarly, we observed increased ALT, AST, BUN, and ferritin in the serum of mice subjected to TNF-α and IFN-γ shock **(Figure 2C)**. Additionally, a meta-analysis of nine studies of patients with COVID-19 showed that thrombocytopenia is associated with a five-fold increased risk of severity and mortality (Lippi et al., 2020). In our murine model, complete blood counts (CBC) revealed a decrease in the number of thrombocytes and percentage of plateletcrit in the blood of mice co-treated with TNF-α and IFN-γ compared with the PBS-treated group **(Figure 2D)**. Although alterations in RBC count and hematocrit percentage have not been reported in patients with COVID-19, we observed increased RBC count, hematocrit percentage, and hemoglobin levels in the blood of mice subjected to TNF-α and IFN-γ shock **(Figure 2D)**.

Lymphopenia is one of the hallmarks of severity and hospitalization in patients with COVID-19 (Tan et al., 2020; Zhao et al., 2020). A meta-analysis revealed an increased neutrophil-to-lymphocyte ratio (NLR) in patients with severe COVID-19 compared with patients with non-severe disease (Chan and Rout, 2020). In our murine model of TNF-α and IFN-γ shock, immune profiling of the peripheral blood identified an overall increase in the relative levels of innate immune cell lineages including macrophages and neutrophils with a reduction in the percentage of B cells and T cells, leading to a concomitant increase in NLR in TNF-α and IFN-γ–treated mice compared with PBS-treated mice **(Figure 2E)**. Overall, these data suggest that TNF-α and IFN-γ cytokine shock in mice mirrors the symptoms of COVID-19.

### TNF-α and IFN-γ induce pyroptosis, apoptosis, and necroptosis

Studies have linked cytokine storm to lung injury, multiple organ failure, and poor prognosis for patients with COVID-19 (Jose and Manuel, 2020; Mehta et al., 2020). Cytokine storm may cause this organ damage through inflammatory cell death. Pyroptosis and necroptosis have been defined as inflammatory cell death pathways. Furthermore, there is crosstalk between these pathways and apoptosis. Therefore, defining thoroughly the nature of cell death induced by the combination of TNF-α and IFN-γ, which had previously been reported to induce immunologically silent apoptosis (Selleri et al., 1995), and dissecting the signaling pathways involved in this process is critical to identify approaches for specific targeting of these molecules.

First, to investigate whether TNF-α and IFN-γ together can induce inflammatory cell death in the form of pyroptosis, we monitored the cleavage of GSDMD. Co-treatment with TNF-α and IFN-γ induced a small amount of cleavage of GSDMD to produce the active P30 fragment that can form membrane pores to induce pyroptosis **(Figure 3A)**. GSDMD can be processed to release this P30 fragment by caspase-1, downstream of inflammasome activation, or caspase-11 (He et al., 2015; Kayagaki et al., 2015; Shi et al., 2015). Consistent with the small amount of GSDMD P30 production, there was no activation of caspase-1 and minimal cleavage of caspase-11 **(Figure 3A)**. Another member of the gasdermin family, GSDME, has also been shown to induce pyroptosis under specific conditions (Wang et al., 2017). We observed that BMDMs stimulated with the combination of TNF-α and IFN-γ displayed robust cleavage of pyroptotic GSDME **(Figure 3A)**, demonstrating that TNF-α and IFN-γ together induced pyroptosis in BMDMs.

**Figure 3.**
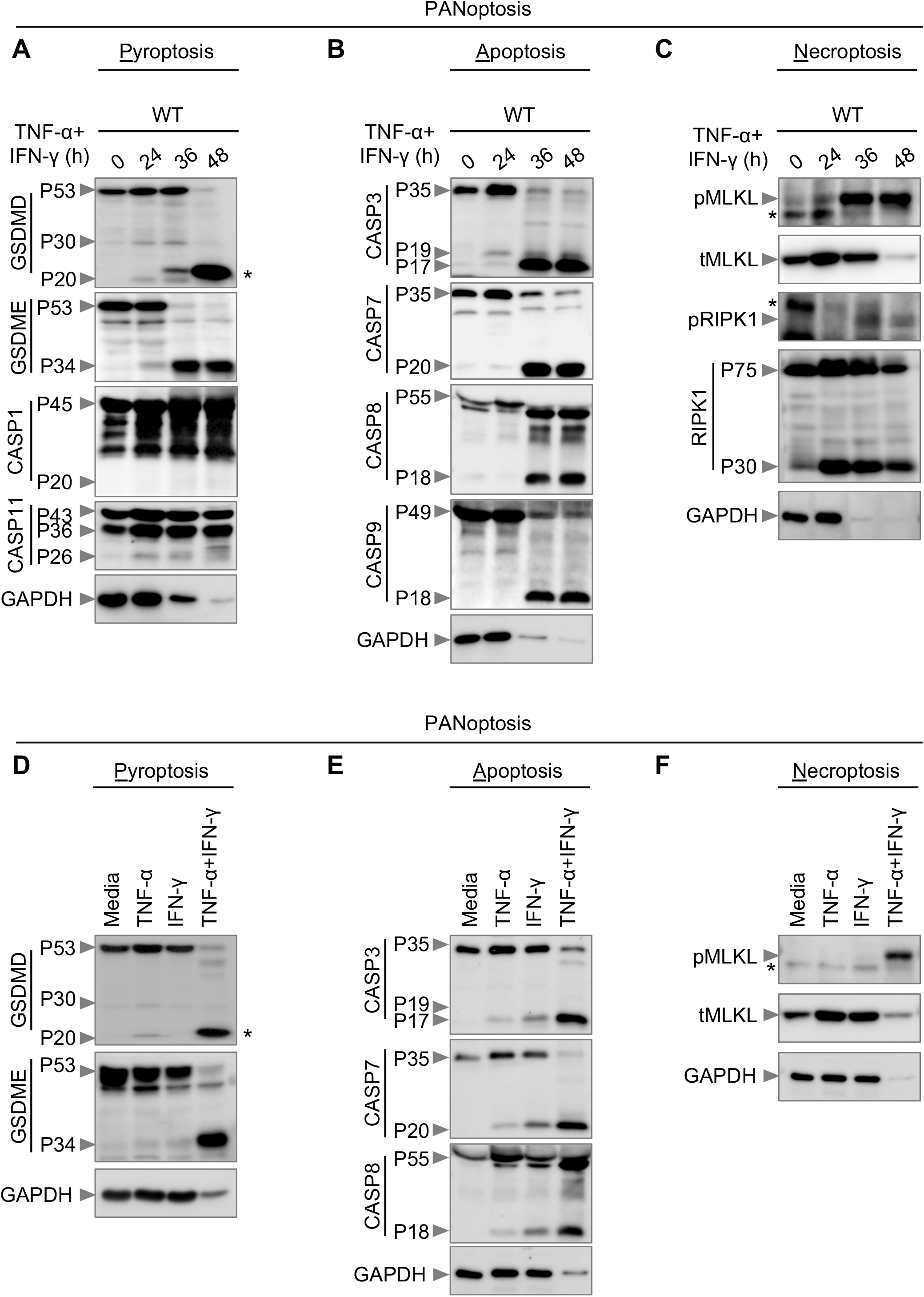
Co-treatment of TNF-α and IFN-γ induces PANoptosis. (A–C) Immunoblot analysis of (A) pro- (P53), activated (P30), and inactivated (P20) GSDMD, pro- (P53) and activated (P34) GSDME, pro- (P45) and activated (P20) CASP1, and pro- (P43) and cleaved CASP-11 (P36 and P26); (B) pro- (P35) and cleaved CASP3 (P19 and P17), pro- (P35) and cleaved CASP-7 (P20), pro- (P55) and cleaved CASP8 (P18), and pro- (P49) and cleaved CASP9 (P18); and (C) phosphorylated MLKL (pMLKL), total MLKL (tMLKL), phosphorylated RIPK1 (pRIPK1), and pro- (P75) and cleaved (P30) RIPK1 in BMDMs co-treated with TNF-α and IFN-γ. (D–F) Immunoblot analysis of (D) GSDMD and GSDME; (E) CASP3, CASP7, and CASP8; and (F) pMLKL and tMLKL in BMDMs after treatment with TNF-α alone, IFN-γ alone, or cotreatment with TNF-α and IFN-γ for 36 h. GAPDH was used as the internal control. Asterisks denote a nonspecific band. Data are representative of at least three independent experiments.

In addition to pyroptosis, we also found that TNF-α and IFN-γ induced apoptosis in BMDMs as evidenced by the cleavage of apoptotic caspases caspase-3, −7, −8, and −9 (**Figure 3B)**. Furthermore, recent studies have shown that activation of caspase-3 and −7 can inactivate GSDMD by processing it to produce a P20 fragment (Chen et al., 2019; Taabazuing et al., 2017), which we also observed **(Figure 3A)**.

Next, we examined whether the combination of TNF-α and IFN-γ induced necroptosis. Cells stimulated with TNF-α and IFN-γ showed robust phosphorylation of MLKL **(Figure 3C)**, suggesting that necroptosis is occurring. During necroptosis, phosphorylation of MLKL occurs downstream of activation of the protein kinases RIPK1 and RIPK3. During TNF-α and IFN-γ treatment, we observed phosphorylation of RIPK1 **(Figure 3C)**. We also noted cleavage of total RIPK1 **(Figure 3C)**, which is involved in regulating apoptosis and necroptosis (Newton et al., 2019).

To investigate the relative contribution of TNF-α or IFN-γ in activating these cell death pathways, we stimulated BMDMs with TNF-α alone, IFN-γ alone, or the combination of TNF-α and IFN-γ. TNF-α or IFN-γ alone did not induce robust cleavage of GSDMD and GSDME **(Figure 3D)**. In addition, we observed reduced cleavage of caspase-3, −7, and −8 in cells stimulated with TNF-α or IFN-γ compared with the combination **(Figure 3E)**. Furthermore, TNF-α or IFN-γ failed to induce robust MLKL phosphorylation **(Figure 3F)**. However, the combination of TNF-α and IFN-γ synergistically induced cleavage of GSDME, caspase-3, −7, and −8, and phosphorylation of MLKL **(Figures 3A–F)**. Collectively, these data suggest that TNF-α and IFN-γ together sensitize the cells to undergo PANoptosis, inflammatory cell death involving the components of pyroptosis, apoptosis, and necroptosis.

### STAT1/IRF1 axis is required for TNF-α and IFN-γ–induced inflammatory cell death

The best-described signaling pathways for TNF-α and IFN-γ are NF-kB and IFN signaling pathways, respectively; however, these pathways have counteracting effects. While NF-kB activation generally drives pro-survival signaling (Liu et al., 2017; Papa et al., 2004), IFN-γ– mediated signaling is cytotoxic (Lin et al., 2017). To understand how these signaling pathways were affected by the combination of TNF-α and IFN-γ, we performed a microarray analysis to identify the most highly upregulated type II IFN-responsive genes in WT BMDMs co-treated with TNF-α and IFN-γ (**Figure 4A)**. In parallel, to determine which genes had relevance in human patients, we re-analyzed a publicly available dataset for differentially regulated type II IFN-responsive genes in patients with differing severities of COVID-19 **(Figure 4B)** (Hadjadj et al., 2020). We found the genes encoding IRF1, IRF5, IRF7, JAK2, and PML to be highly upregulated in both patients with severe COVID-19 and TNF-α and IFN-γ–treated BMDMs. To investigate the role of IRF1, IRF5, and IRF7 in instigating inflammatory cell death in response to TNF-α and IFN-γ, *Irf1^-/-^, Irf5^-/-^* and *Irf7^-/-^* BMDMs were analyzed. Cells deficient in IRF1, but not IRF5 or IRF7, were protected from cell death upon treatment with TNF-α and IFN-γ **(Figures 4C and S3A)**. JAK2 is known to signal upstream of IRF1 through STAT1; autophosphorylation of JAK2 phosphorylates JAK1 to activate the transcription factor STAT1, which localizes to the nucleus to induce transcription of type II IFN-responsive genes, including IRF1 (Schroder et al., 2004). Based on the upregulation of JAK2 in both patients with severe COVID-19 and BMDMs treated with TNF-α and IFN-γ **(Figures 4A and 4B)** and based on the confirmed role of IRF1 in promoting cell death in this pathway **(Figure 4C)**, we also assessed the role of upstream molecules in TNF-α and IFN-γ–mediated cell death. Similar to IRF1-deficient cells, BMDMs lacking STAT1 were protected from cell death **(Figures S3A–S3C)**. Consistent with this protection, *Irf1^-/-^* BMDMs showed impaired activation of apoptotic caspases (caspase-3, −7, and −8); the pyroptotic molecule GSDME; and the necroptotic molecule MLKL **(Figures S3D and S3E)**. It is possible that type II IFN-responsive genes other than IRF1 that were induced in patients with severe COVID-19 or TNF-α and IFN-γ co-treated BMDMs also contributed to the cell death. To determine whether these other molecules were contributing, we looked at the cell death in cells lacking molecules involved in IFN production or signaling including IRF1, IRF2, IRF3, IRF5, IRF7, IFNAR1, IFNAR2, IRF9, STAT1, TRIF, MDA5, MAVS, cGAS, STING, GBP2, and PTPN6. We observed that TNF-α and IFN-γ co-treatment– induced cell death was only impaired in IRF1- and STAT1-deficient BMDMs **(Figure S3A**). Altogether, these findings suggest that the STAT1/IRF1 axis regulates inflammatory cell death in response to TNF-α and IFN-γ treatment.

**Figure 4.**
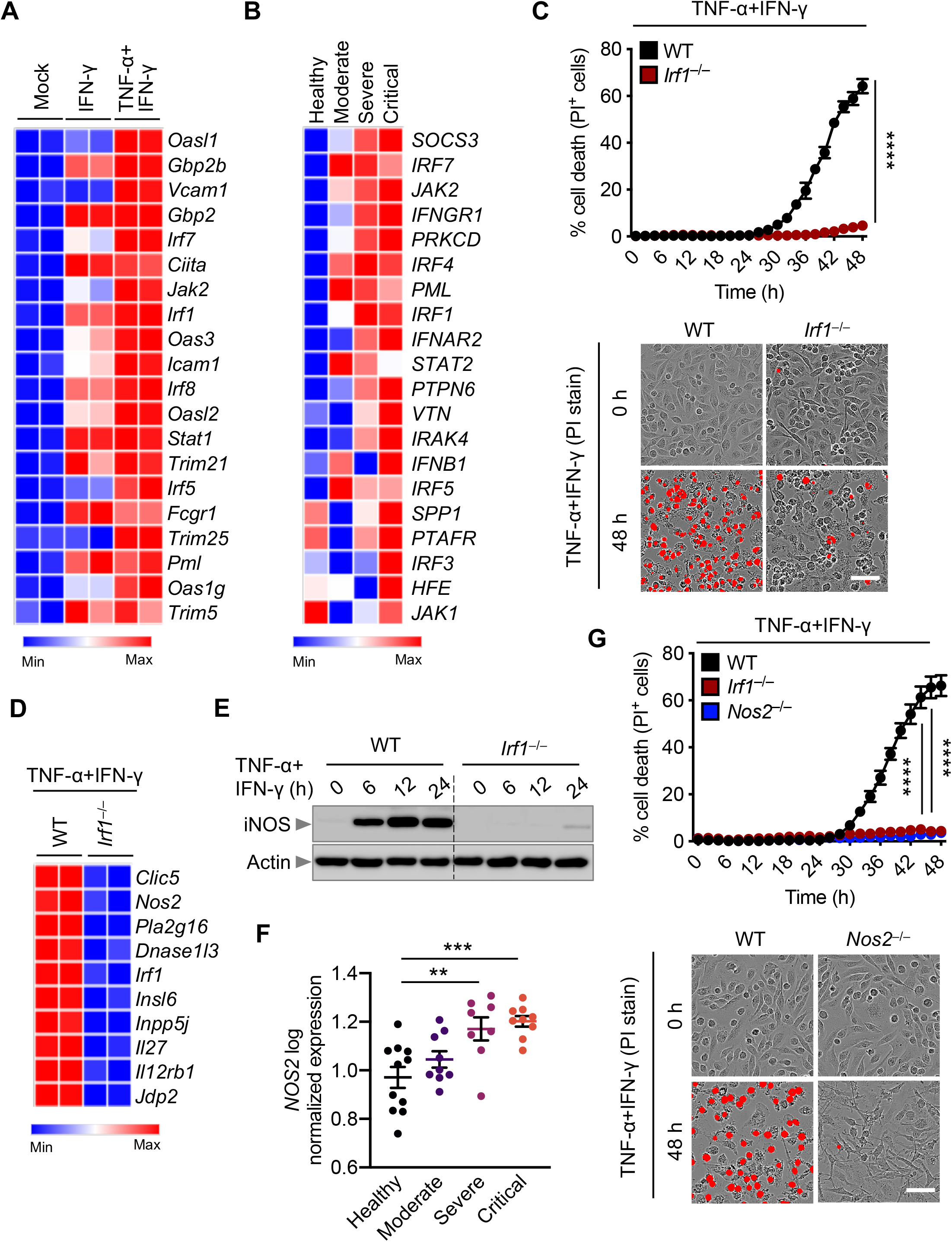
IRF1 and NOS2 mediate TNF-α and IFN-γ–induced inflammatory cell death. (A) Heatmap depicting the expression levels of type II IFN-responsive genes in WT BMDMs treated with IFN-γ alone or co-treated with TNF-α and IFN-γ for 16 h relative to their expression in untreated (Mock) BMDMs. (B) Heatmap depicting the expression levels of type II IFN-responsive genes in patients with moderate, severe, and critical COVID-19 relative to their expression in healthy patients (Hadjadj et al., 2020). (C) Real-time analysis of cell death in TNF-α and IFN-γ co-treated WT and *Irf1^-/-^* BMDMs. Representative images of cell death are shown at 0 h and after 48 h of TNF-α and IFN-γ treatment. (D) Heatmap depicting the expression levels of the most downregulated genes in *Irf1^-/-^* BMDMs co-treated with TNF-α and IFN-γ for 16 h relative to their expression in WT treated BMDMs. (E) Immunoblot analysis of iNOS in WT and *Irf1^-/-^* BMDMs co-treated with TNF-α and IFN-γ. Actin was used as the internal control. (F) Expression analysis of *NOS2* in patients with moderate, severe, and critical COVID-19 relative to the expression in healthy patients (Hadjadj et al., 2020). (G) Real-time analysis of cell death in WT, *Irf1^-/-^*, and *Nos2^-/-^* BMDMs during co-treatment with TNF-α and IFN-γ. Representative images of cell death are shown at 0 h and after 48 h of TNF-α and IFN-γ treatment. Scale bar, 50 μm. Data are representative of at least three independent experiments. \*\**P* < 0.01; \*\*\**P* < 0.001; \*\*\*\**P* < 0.0001. Analysis was performed using the one-way ANOVA (F) or two-way ANOVA (C and G). Data are shown as mean ± SEM (C, F, and G). See also Figure S3–S5.

Given that IRF1 is a transcription factor, it is likely that it regulates inflammatory cell death by inducing type II IFN-responsive genes. Using a microarray, we identified genes which had the lowest levels of expression in *Irf1^-/-^* BMDMs co-stimulated with TNF-α and IFN-γ compared with the similarly stimulated WT BMDMs. We identified *Nos2*, the gene encoding inducible nitric oxide synthase (iNOS), as one of the most downregulated genes in *Irf1^-/-^* BMDMs under these conditions **(Figure 4D)**. The protein levels of iNOS were also reduced in *Irf1^-/-^* cells treated with TNF-α and IFN-γ **(Figure 4E)**. In addition, iNOS protein expression was abolished in *Stat1^-/-^* cells during treatment with TNF-α and IFN-γ **(Figure S4A)**. Consistent with the reduced iNOS expression, we observed reduced production of nitric oxide (NO) in *Irf1^-/-^* and *Stat1^-/-^* cells compared with WT cells upon treatment with TNF-α and IFN-γ **(Figure S4B)**. Together, these results suggest that iNOS is an IFN-inducible protein which is produced through a mechanism requiring STAT1 and IRF1 downstream of TNF-α and IFN-γ stimulation.

To determine whether iNOS had any impact in COVID-19 pathogenesis, we analyzed a publicly available dataset and found that *NOS2* is significantly upregulated in patients with severe and critical COVID-19 compared with healthy controls **(Figure 4F)** (Hadjadj et al., 2020). iNOS and the downstream NO are involved in numerous biological processes (Xu et al., 2002), and NO can be cytotoxic or cytostatic depending upon the context (Albina and Reichner, 1998; Pervin et al., 2001). To investigate whether iNOS and NO play important roles in the cell death induced by TNF-α and IFN-γ treatment, we assessed *Nos2^-/-^* BMDMs, which lack iNOS expression **(Figure S4C)**. Upon treatment with TNF-α and IFN-γ, *Nos2^-/-^* BMDMs were protected from cell death **(Figure 4G)**. We also treated WT BMDMs with NO production inhibitors and found that both L-NAME and 1400W inhibited cell death in response to TNF-α and IFN-γ treatment **(Figures S4D– S4F)**. Consistent with the impaired cell death, *Nos2^-/-^* BMDMs had reduced activation of apoptotic caspases (caspase-3, −7, and −8), the pyroptotic molecule GSDME, and the necroptotic molecule MLKL **(Figures S4G and S4H)**. Overall, these data suggest that the STAT1/IRF1 axis regulates iNOS expression for the production of NO, which subsequently induces inflammatory cell death in response to TNF-α and IFN-γ co-treatment.

While TNF-α or IFN-γ alone have been shown to induce NO (Salim et al., 2016), they each failed to induce cell death on their own **(Figures 1C and 1E)**. Moreover, NO has been demonstrated to be cytotoxic or cytostatic depending upon the context (Albina and Reichner, 1998; Pervin et al., 2001). Therefore, it is possible that the concentration of NO is critical to induce cell death and that synergism of TNF-α and IFN-γ results in a concomitant increase in the concentration of NO to reach the threshold necessary to activate cell death pathways. Indeed, TNF-α and IFN-γ synergistically induced iNOS and produced more NO than TNF-α or IFN-γ alone **(Figures S5A and S5B)**. To further confirm that the cytotoxicity of NO was concentration-dependent, we treated WT BMDMs with increasing concentrations of the NO donor SIN-1. We observed that the kinetics of cell death induced by SIN-1 were proportional to the NO concentration **(Figure S5C)**, suggesting that the level of NO produced by either TNF-α or IFN-γ alone may not be sufficient to induce cell death. Altogether, these data indicate that TNF-α and IFN-γ synergistically induce iNOS and NO, which subsequently induce cell death.

It is also possible that cell death in response to TNF-α and IFN-γ may be induced through an additional route; IFN-γ may interfere with TNF-α signaling to suppress NF-κB activation and switch the pro-survival signaling of TNF-α to cell death. TNF-α signaling engages RIPK1 to induce caspase-8–driven apoptosis or RIPK3-driven necroptosis in the context of NF-κB inhibition or caspase-8 inhibition, respectively (Lin et al., 1999; Vercammen et al., 1998). To address the possibility that IFN-γ is suppressing NF-κB to drive TNF-α–induced cell death, we re-analyzed our microarray data to assess the expression of NF-κB target genes for inflammatory cytokines/chemokines and apoptosis regulators in WT BMDMs treated with TNF-α alone or the combination of TNF-α and IFN-γ. We found that the expression NF-κB target genes for inflammatory cytokines/chemokines was not impaired in the cells stimulated with TNF-α and IFN-γ compared with their expression in cells stimulated with TNF-α alone **(Figure S5D)**. Moreover, treatment with IFN-γ did not influence TNF-α–driven expression of genes that regulate apoptosis **(Figure S5E)**. These data indicate that IFN-γ does not suppress TNF-α–induced NF-kB signaling to induce cell death. Collectively, our data show that the cell death induced by TNF-α and IFN-γ is driven by the IRF1/STAT1 pathway and involves the induction of iNOS and NO, which subsequently cause cell death.

### RIPK1/FADD/CASP8 axis drives PANoptosis induced by TNF-α and IFN-γ

NO is known to induce apoptosis by activating caspase-8–dependent pathways (Du et al., 2006; Dubey et al., 2016). In addition to its classical role in inducing apoptosis, caspase-8 plays a critical role in regulating inflammatory cell death pathways, pyroptosis and necroptosis (Fritsch et al., 2019; Gurung et al., 2014; Gurung et al., 2016; Kuriakose et al., 2016; Lukens et al., 2014; Malireddi et al., 2020; Malireddi et al., 2018; Newton et al., 2019; Orning et al., 2018; Sarhan et al., 2018). To determine whether caspase-8 mediates the cell death induced by TNF-α and IFN-γ, we analyzed BMDMs derived from *Ripk3^-/-^* and *Ripk3^-/-^ Casp8^-/-^* mice, since caspase-8– deficient mice are not viable. RIPK3 deficiency failed to protect against cell death, whereas deletion of both RIPK3 and caspase-8 provided substantial protection against the cell death induced by TNF-α and IFN-γ co-treatment **(Figures 5A and 5B)**. The importance of caspase-8 in inflammatory cell death in response to TNF-α and IFN-γ was further supported by impaired activation of apoptotic caspases (caspase-3 and −7); the pyroptotic molecule GSDME; and the necroptotic molecule MLKL in *Ripk3^-/-^ Casp8^-/-^* cells compared with their activation in WT cells **(Figures 5C and 5D)**.

**Figure 5.**
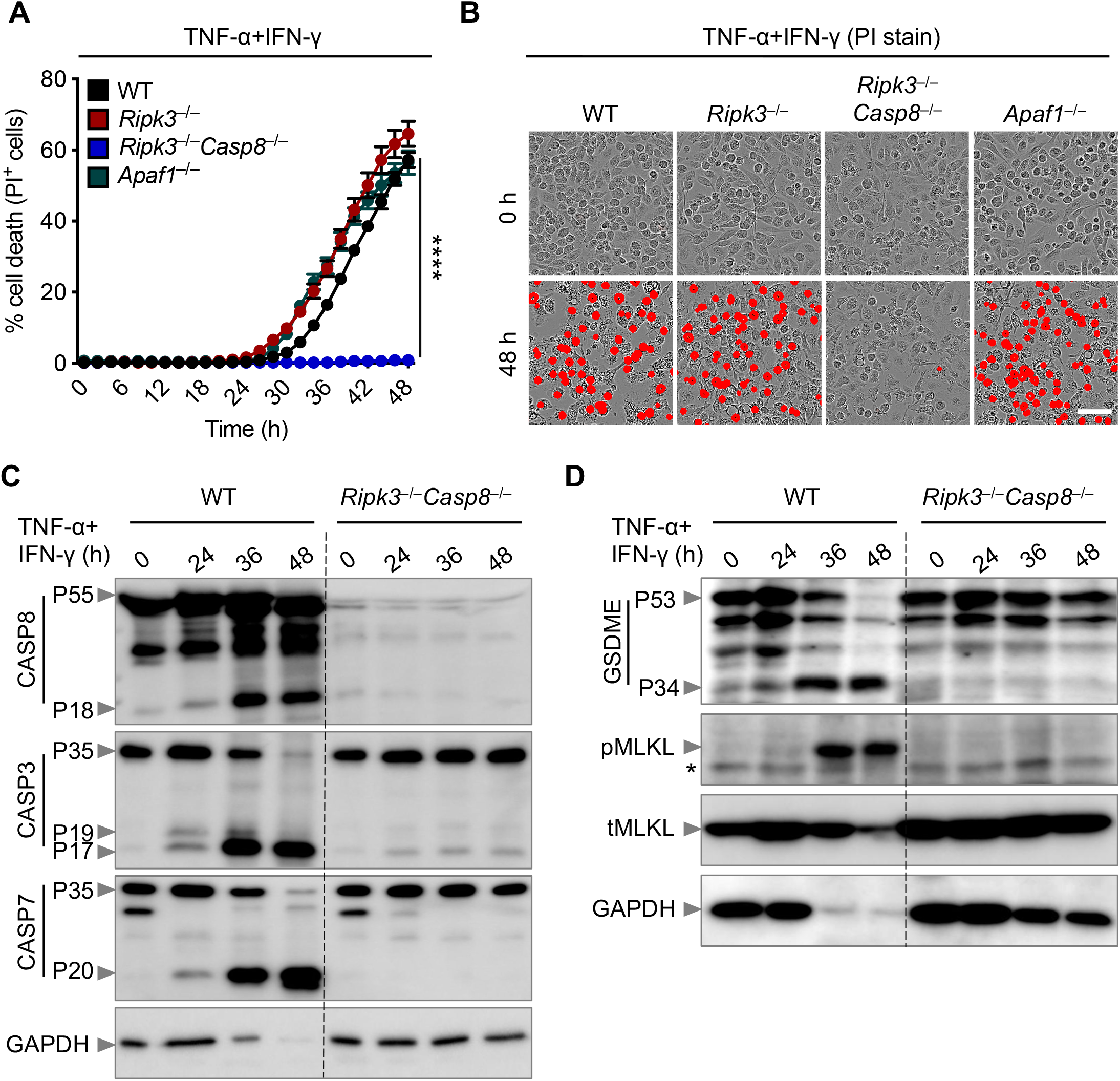
Caspase-8 drives PANoptosis induced by co-treatment with TNF-α and IFN-γ. (A) Real-time analysis of cell death in WT, *Ripk3^-/-^, Ripk3^-/-^ Casp8^-/-^*, and *Apaf1^-/-^* BMDMs cotreated with TNF-α and IFN-γ. (B) Representative images of cell death in WT, *Ripk3^-/-^, Ripk3^-/-^ Casp8^-/-^*, and *Apaf1^-/-^* BMDMs. Scale bar, 50 μm. (C, D) Immunoblot analysis of (C) pro- (P55) and cleaved CASP8 (P18), pro- (P35) and cleaved CASP3 (P19 and P17), and pro- (P35) and cleaved CASP-7 (P20); and (D) pro- (P53) and activated (P34) GSDME, phosphorylated MLKL (pMLKL), and total MLKL (tMLKL) in WT and *Ripk3^-/-^ Casp8^-/-^* BMDMs co-treated with TNF-α and IFN-γ. GAPDH was used as the internal control. Asterisks denote a nonspecific band. Data are representative of at least three independent experiments. \*\*\*\**P* < 0.0001. Analysis was performed using the two-way ANOVA. Data are shown as mean ± SEM (A). See also Figures S6 and S7.

Caspase-8 is recruited to cell death-inducing complexes by FADD. To address whether FADD regulates pyroptosis, apoptosis, and necroptosis induced by TNF-α and IFN-γ co-treatment, we examined BMDMs derived from *Ripk3^-/-^* and *Ripk3^-/-^ Fadd^-/-^* mice, since FADD-deficient mice are not viable. As before, loss of RIPK3 did not protect cells from undergoing death; however, deletion of both RIPK3 and FADD provided substantial protection against the cell death induced by TNF-α and IFN-γ co-treatment **(Figures S6A and S6B)**. We also observed reduced activation of caspase-3, −7, and −8; GSDME; and MLKL in *Ripk3^-/-^ Fadd^-/-^* cells compared with WT cells **(Figures S6C and S6D)**. These data suggest that caspase-8 and FADD are key regulators of PANoptosis in response to TNF-α and IFN-γ.

APAF1/caspase-9 are known to be mediators of the intrinsic apoptotic pathway. To study the role of intrinsic apoptosis in response to TNF-α and IFN-γ stimulation, we used BMDMs deficient in APAF1 and found similar dynamics of cell death between WT and *Apaf1^-/-^* BMDMs **(Figures 5A and 5B)**, suggesting that intrinsic apoptosis does not regulate the cell death triggered by TNF-α and IFN-γ co-treatment in macrophages. Collectively, these data suggest that the FADD/caspase-8 axis regulates TNF-α and IFN-γ co-treatment–induced inflammatory cell death independent of intrinsic apoptosis in macrophages.

Next, we determined the contribution of downstream molecules activated by caspase-8 in TNF-α and IFN-γ–induced cell death. BMDMs deficient in caspase-3, but not caspase-7, showed reduced cell death with respect to WT cells **(Figure S7A)**. Loss of the necroptotic executioner MLKL or the pyroptotic executioner GSDMD failed to protect cells against death triggered by TNF-α and IFN-γ co-treatment **(Figure S7B)**. In addition, loss of the upstream activators of GSDMD, caspase-1 and caspase-11, did not protect against cell death **(Figure S7C)**. Similar to caspase-3–deficient cells, cells lacking GSDME showed reduced cell death compared with WT cells in response to TNF-α and IFN-γ co-treatment **(Figure S7B)**. However, combined deletion of GSDME with the other pore-forming molecules MLKL and GSDMD showed a similar level of protection as that of cells lacking GSDME alone **(Figure S7B)**, suggesting that GSDME and caspase-3 potentiate the inflammatory cell death in response to TNF-α and IFN-γ co-treatment.

### Inhibition of inflammatory cell death, PANoptosis, provides protection against lethality induced by TNF-α and IFN-γ

Based on our findings that the STAT1/IRF1/caspase-8 axis drives inflammatory cell death downstream of TNF-α and IFN-γ, we hypothesized that inhibition of this pathway would provide protection against TNF-α and IFN-γ–induced shock *in vivo*. Indeed, mice deficient in STAT1 or both RIPK3 and caspase-8 were resistant to mortality induced by the cytokine shock, while mice deficient in RIPK3 alone were not **(Figures 6A and 6B)**, suggesting that STAT1-mediated, caspase-8–dependent cell death contributes to the fatal outcome. Next, we investigated whether deletion of STAT1 or caspase-8 would rescue the laboratory abnormalities caused by TNF-α and IFN-γ shock that are consistent with the symptoms observed in patients with COVID-19. There was increased cell death as measured by serum LDH in WT mice subjected to TNF-α and IFN-γ shock compared with PBS–treated WT mice **(Figures 2C and 6C)**. Consistent with the reduced cell death observed *in vitro*, the LDH level in TNF-α and IFN-γ–treated *Stat1^-/-^* or *Ripk3^-/-^ Casp8^-/-^* mice was similar to that in PBS-treated *Stat1^-/-^, Ripk3^-/-^ Casp8^-/-^*, or WT mice **(Figure 6C)**, suggesting that STAT1– and caspase-8–mediated cell death is driving lethality in mice subjected to TNF-α and IFN-γ shock. Similarly, deletion of STAT1 or caspase-8 reduced the level of ALT and AST in mice treated with TNF-α and IFN-γ shock **(Figure 6C)**. However, *Stat1^-/-^* and *Ripk3^-/-^ Casp8^-/-^* mice co-treated with TNF-α and IFN-γ had a decreased percentage of T cells in the blood when compared to their PBS-treated counterparts **(Figure 6D)**. The number of thrombocytes and percentage of plateletcrit in the blood of *Stat1^-/-^* or *Ripk3^-/-^ Casp8^-/-^* mice cotreated with TNF-α and IFN-γ were generally similar to those of WT mice treated with PBS **(Figure 6E)**. Moreover, RBC count, hematocrit percentage, and hemoglobin levels in the blood of TNF-α and IFN-γ–treated *Stat1^-/-^* or *Ripk3^-/-^ Casp8^-/-^* mice were similar to those in PBS-treated *Stat1^-/-^, Ripk3^-/-^ Casp8^-/-^*, or WT mice **(Figure 6F)**. Altogether, these data suggest that inhibition of inflammatory cell death driven by the STAT1/caspase-8 axis is crucial to prevent TNF-α and IFN-γ–mediated pathology and mortality *in vivo*.

**Figure 6.**
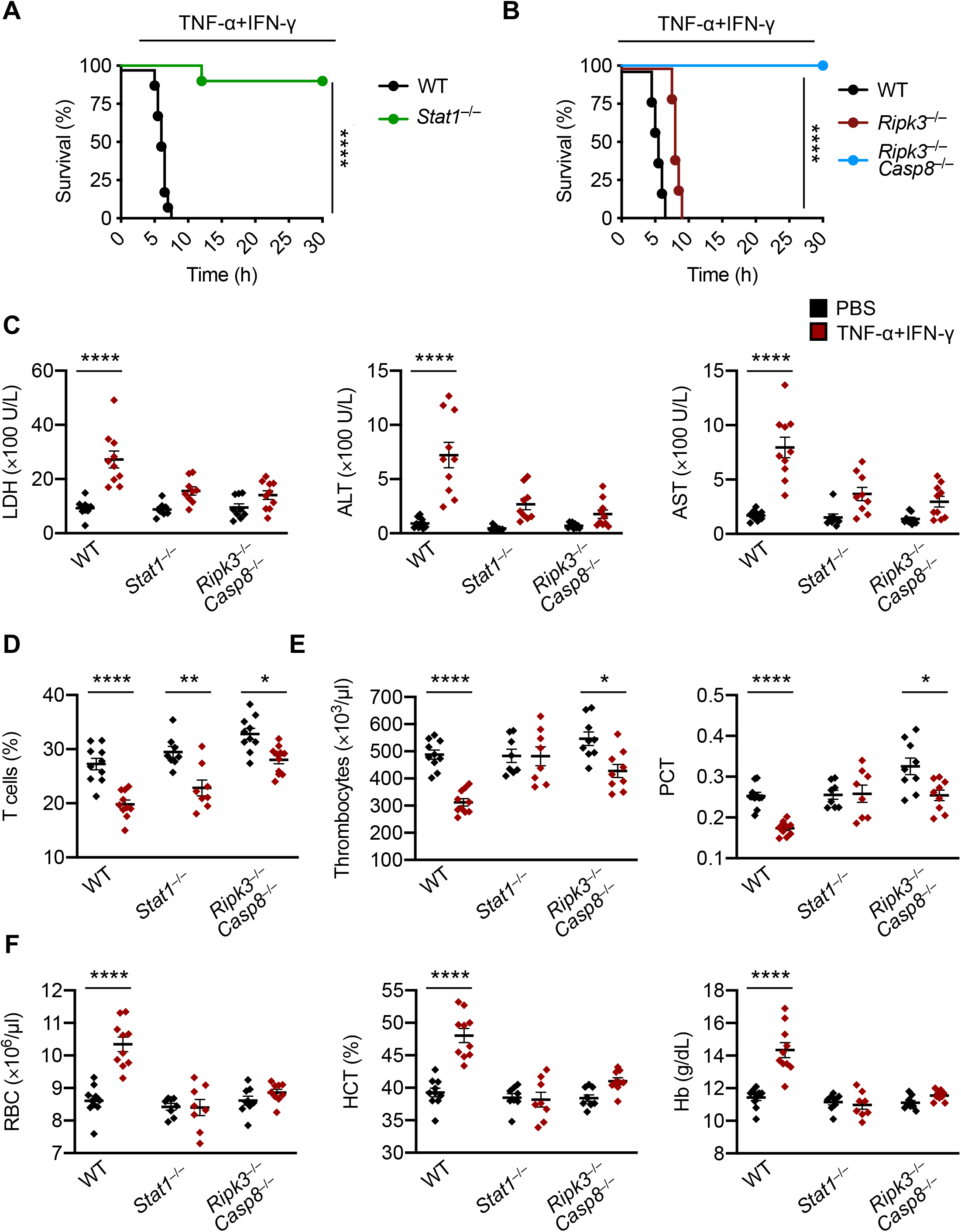
Inhibition of PANoptosis provides protection against TNF-α and IFN-γ–driven lethality in mice. (A, B) Survival of 6-to 8-week-old (A) WT (n = 10) and *Stat1^-/-^* (n = 10) mice; and (B) WT (n = 10), *Ripk3^-/-^* (n = 12), and *Ripk3^-/-^ Casp8^-/-^* (n = 15) mice after i.p. injection of TNF-α and IFN-γ. (C–E) Analysis of (C) serum levels of LDH, ALT, and AST; (D) percentage of T cells in blood; (E) the number of thrombocytes and plateletcrit (PCT) in the blood; and (F) RBC count, hematocrit (HCT), and hemoglobin (Hb) concentration in the blood of WT, *Stat1^-/-^*, and *Ripk3^-/-^ Casp8^-/-^* mice injected intraperitoneally with PBS or TNF-α and IFN-γ at 5 h post-treatment. Data are representative of two independent experiments. Data are shown as mean ± SEM (C–F). \**P* < 0.05; \*\**P* < 0.01; \*\*\*\**P* < 0.0001. Analysis was performed using the survival curve comparison (log-Rank [Mantel-Cox] test) (A and B) or the two-way ANOVA (C–F).

### Blocking TNF-α and IFN-γ reduces mortality in disease models associated with cytokine storm: SARS-CoV-2 infection, cytokine shock, sepsis, and HLH

Our *in vitro* and *in vivo* studies show that the synergistic activity of the pro-inflammatory cytokines TNF-α and IFN-γ mimic the symptoms of COVID-19 in mice and trigger robust cell death. These findings suggest that blocking TNF-α and IFN-γ during SARS-CoV-2 infection would prevent severe symptoms and protect against death. To understand the potential impact of blocking TNF-α and IFN-γ during disease, we tested the efficacy of treatment with neutralizing antibodies against TNF-α and IFN-γ. Mice injected with the neutralizing antibodies, but not isotype control, were 100% protected from death during TNF-α and IFN-γ shock **(Figure 7A)**, indicating that these antibodies efficiently neutralized TNF-α and IFN-γ *in vivo*. To test the potential efficacy of this treatment during SARS-CoV-2 infection, we used a murine SARS-CoV-2 infection model (Winkler et al., 2020). By 7 days post-infection, nearly all isotype control-treated mice succumbed to the infection. Conversely, treatment with neutralizing antibodies against TNF-α and IFN-γ provided significant protection against SARS-CoV-2–induced mortality **(Figure 7B)**.

**Figure 7.**
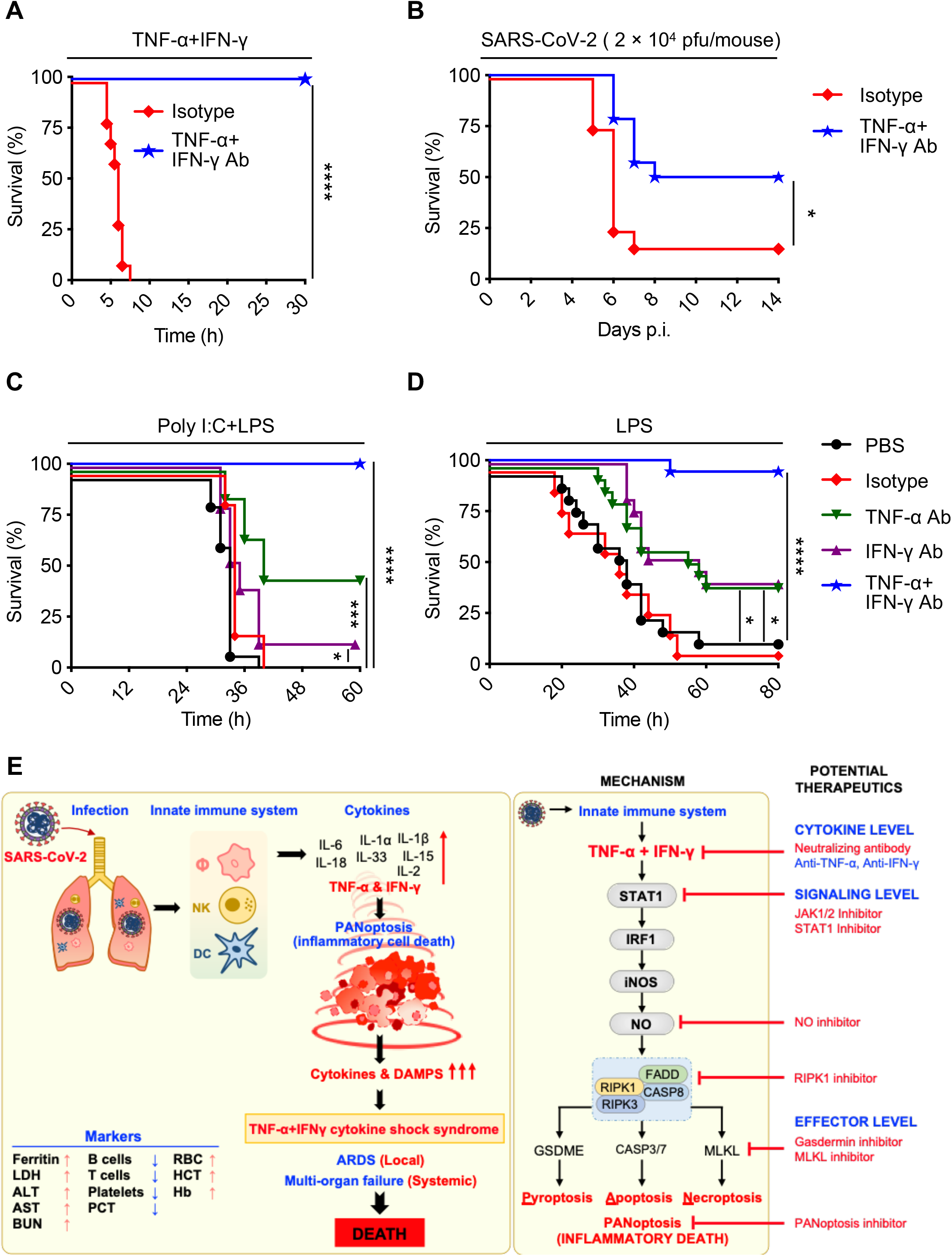
Blocking TNF-α and IFN-γ provides protection in disease models associated with cytokine storm: SARS-CoV-2 infection, cytokine shock, sepsis, and HLH. (A) Survival of 6-to 8-week-old WT mice injected with isotype control (n = 10) or neutralizing antibodies against TNF-α and IFN-γ (n = 14) 12 h before i.p. injection of TNF-α and IFN-γ. (B) Survival of 7-to 8-week-old K18-hACE2 transgenic mice injected with isotype control (n = 12) or neutralizing antibodies against TNF-α and IFN-γ (n = 13) on day 1, 3, and 4 after infection with SARS-CoV-2 (2 × 10^4^ pfu/mouse). (C) Survival of 7-to 8-week-old WT mice injected intraperitoneally with poly I:C (10 mg/kg body weight) followed 24 h later by i.p. injection of PBS (n = 15), isotype control (n = 15), neutralizing antibody against TNF-α (n = 15), neutralizing antibody against IFN-γ (n = 15), or neutralizing antibodies against both TNF-α and IFN-γ (n = 15). Mice were then challenged with LPS (5 mg/kg body weight) 1 h after the treatments. (D) Survival of 7-to 8-week-old WT mice injected with PBS (n = 17), isotype control (n = 10), neutralizing antibody against TNF-α (n = 17), neutralizing antibody against IFN-γ (n = 17), or neutralizing antibodies against both TNF-α and IFN-γ (n = 18) 30 min and 6 h after intraperitoneal injection of a lethal dose of LPS (20 mg/kg body weight). (E) Schematic overview of the mechanism of TNF-α and IFN-γ–induced pathology and the inflammatory cell death pathway with strategies for potential therapeutics. Data are pooled from two independent experiments (A–D). \**P* < 0.05; \*\*\**P* < 0.001; \*\*\*\**P* < 0.0001. Analysis was performed using the survival curve comparison (log-Rank [Mantel-Cox] test).

To determine the potential therapeutic benefit of blocking this pathway in other diseases and syndromes associated with cytokine storm, we tested the efficacy of neutralizing antibodies against TNF-α and IFN-γ in other models. Poly I:C priming followed by LPS challenge recapitulates most aspects of HLH, a severe systemic inflammatory syndrome associated with life-threatening symptoms (Wang et al., 2019). Treatment with the combination of TNF-α and IFN-γ blocking antibodies provided 100% protection against the lethality induced by poly I:C and LPS challenge **(Figure 7C)**. Next we investigated the therapeutic potential of these neutralizing antibodies in a murine sepsis model. Injection of a lethal dose of LPS induced mortality in 90% of mice by 48 h. Neutralization of TNF-α alone or IFN-γ alone provided some protection, but combined neutralization of TNF-α and IFN-γ provided 94% protection against LPS-induced lethality **(Figure 7D)**. These *in vivo* models suggest that TNF-α and IFN-γ–mediated inflammatory cell death, PANoptosis, drives pathology in not only COVID-19, but also other diseases and syndromes associated with cytokine storm, including cytokine shock, sepsis, and HLH.

## DISCUSSION

Fatalities in patients with severe COVID-19 are linked to elevated levels of circulating pro-inflammatory cytokines, or cytokine storm (Jose and Manuel, 2020; Mehta et al., 2020; Ragab et al., 2020). Virtually every cell in the body is capable of responding to cytokines. Some pro-inflammatory cytokines such as IL-1β, TNF-α, and IFN-γ induce cell death in various cell types, and this cell death is linked to pathological conditions such as neurological disorders, liver damage, COPD, osteoporosis, HLH, sepsis, and more (Belkhelfa et al., 2014; De Boer, 2002; Sass et al., 2001; Yang et al., 2020; Zheng et al., 2016). However, the cellular and molecular mechanisms employed by these cytokines in inducing cytokine storm and inflammatory cell death have not been well described. Our extensive characterization of COVID-19 cytokines here showed that TNF-α and IFN-γ synergism induces inflammatory cell death, PANoptosis, that is dependent on the IRF1 and NO axis.

The phrase “cytokine storm” has been used as a descriptive term to encompass a range of observations that are linked to hyperactivation of the immune system and uncontrolled and excessive release of cytokines which may result in multisystem organ failure and death. While the identity of the cytokines that are released during cytokine storm has been characterized in many diseases, the downstream effects of these cytokines and the pathways propagating the subsequent inflammation are less well understood. Because cytokine storm has been described by many as a hallmark of COVID-19 pathology, several clinical trials are ongoing to assess the efficacy of cytokine blockade using inhibitors of IL-6, IL-18, IL-1α, and IL-1β (Aouba et al., 2020; Atal and Fatima, 2020; Huet et al., 2020; Lythgoe and Middleton, 2020). However, the mechanisms of action of these anti-cytokine therapies in COVID-19 are largely unknown, and there are no systematic studies or genetic evidence to serve as a basis to support the utility of these therapies. Additionally, blocking pro-inflammatory cytokines such as TNF-α, IL-1, or Il-6 have had mixed success in the treatment of other diseases associated with cytokine storm, highlighting the lack of mechanistic understanding of the pathogenic process that accompanies cytokine storm. Furthermore, some have questioned whether the phenomenon observed during COVID-19 truly constitutes a cytokine storm, as emerging evidence has shown that the levels of pro-inflammatory cytokines in these patients are much lower than in patients with other diseases previously associated with cytokine storm, such as sepsis (Kox et al., 2020; Leisman et al., 2020). Our study suggests a new paradigm for defining the mechanism of cytokine storm by focusing on the specific effects of the component cytokines.

Of the cytokines induced in COVID-19, our findings show that TNF-α and IFN-γ play a prominent role in damaging vital organs by inducing inflammatory cell death. Indeed, blocking IL-6 has had mixed results clinically, and our results suggest that blocking cytokines other than TNF-α and IFN-γ may fail as well. In line with the lack of consistent efficacy of IL-6 blockade clinically, we did not see any effect of IL-6 individually or in combination in driving inflammatory cell death *in vitro*. Our *in vivo* findings show that symptoms of TNF-α and IFN-γ shock mirror those of severe SARS-CoV-2 infection in patients, and treatment with the combination of TNF-α and IFN-γ neutralizing antibodies provided protection against SARS-CoV-2 infection in the mouse model. Although the monotherapy with anti-TNF-α or anti-IFN-γ showed minimal protection in non-infection related cytokine storm models, the relative contribution of monotherapy with anti-TNF-α or anti-IFN-γ in providing protection against SARS-CoV-2 infection requires further investigation. Blocking these cytokines together or individually might be useful in mitigating the cytokine storm and preventing severe disease clinically. Case studies have shown that patients who are on anti–TNF-α therapy for autoinflammatory diseases, such as IBD or Crohn’s, who contract SARS-CoV-2 tend to have a mild disease course (Rodriguez-Lago et al., 2020; Waggershauser et al., 2020), suggesting that prophylactic anti-cytokine therapy may also be beneficial. While inhibiting cell death in the context of viral infection has a risk of increasing the viral production from infected cells, a growing body of evidence suggests that excessive cell death contributes more to disease pathology than does viral titer. Studies suggest that exaggerated inflammatory responses, loss of pulmonary epithelia, and acute lung injury, which are all associated with cell death, are the major factors contributing to morbidity and mortality during pathogenic influenza infection (de Jong et al., 2006). Additionally, mice lacking both TNF-α and IL-1 receptors exhibit reduced lung inflammation and a delay in mortality following infection with a highly virulent H5N1 virus (Perrone et al., 2010). Moreover, children tend not to develop severe COVID-19 despite having high viral titers (Kam et al., 2020), and the peak viral titers in respiratory tract samples might occur even before the symptom onset of pneumonia in SARS-CoV and SARS-CoV-2 infections (Peiris et al., 2003). Combined with the protection we observed *in vivo* during SARS-CoV-2 infection conferred by neutralizing TNF-α and IFN-γ, these findings suggest inhibition of TNF-α and IFN-γ signaling might be beneficial and not lead to super-infection with the virus.

In addition to the potential therapeutic impact of blocking TNF-α and IFN-γ, our study delineated the molecular pathway engaged by these cytokines to induce cell death, highlighting a number of additional potential therapeutic targets **(Figure 7E)**. Downstream of TNF-α and IFN-γ, we identified a critical role for STAT1 in promoting inflammatory cell death. Several inhibitors of STAT1 and its upstream regulator JAK are in clinical trials for various diseases (Damsky and King, 2017; Miklossy et al., 2013; Qureshy, 2020). The JAK/STAT1 pathway controls the transcriptional regulation of IRF1, which we also identified as critical for inflammatory cell death in response to TNF-α and IFN-γ. IRF1 has previously been shown to regulate cell death in other studies, including the induction of PANoptosis to suppress colorectal tumorigenesis (Benaoudia et al., 2019; Karki et al., 2020; Kayagaki et al., 2019; Kuriakose et al., 2018; Man et al., 2015). Downstream of IRF1, we identified iNOS and NO as being important for inflammatory cell death. NO can inhibit NLRP3 inflammasome assembly (Hernandez-Cuellar et al., 2012), possibly explaining the inability of TNF-α and IFN-γ to induce caspase-1 and GSDMD activation in our study. However, IFN-γ priming potentiates caspase-11–driven GSDMD cleavage in response to cytosolic LPS (Brubaker et al., 2020). Multiple studies have demonstrated a context-dependent role for NO as a cytotoxic or cytostatic molecule (Albina and Reichner, 1998; Pervin et al., 2001), and clinical trials examining the impact of NO inhibitors have been pursued previously (Bailey et al., 2007; Petros et al., 1991; Wong and Lerner, 2015). Additionally, the potential utility of inhaled NO is currently being evaluated in COVID-19 (Lei et al., 2020). Such treatment would preferentially vasodilate the pulmonary arterioles and improve oxygenation; however, our data suggest that increasing the concentration of NO in the serum is likely to be deleterious. A mathematical modeling to dissect out the roles of each cytokine found that TNF-α is largely responsible for the timing of iNOS induction by inducing a rapid response, whereas IFN-γ contributes to the concentration of NO produced (Salim et al., 2016). Inhibition of executioners of PANoptosis might represent another therapeutic approach to prevent the cell death induced by TNF-α and IFN-γ. While it would be ideal to target these common steps downstream of TNF-α and IFN-γ in the cell death pathway, directly targeting the cytokines together or individually represents the most immediate therapeutic strategy to pursue, as anti–TNF-α and anti–IFN-γ antibodies are already approved for clinical use.

One of the hallmarks of TNF-α and IFN-γ shock or severe COVID-19 is lymphopenia (Laing et al., 2020). NO has been shown to induce T-cell death (Moulian et al., 2001), and T cells lacking iNOS have reduced post-activation death (Vig et al., 2004). Also, T cells lacking STAT1 are resistant to activation-induced cell death (Refaeli et al., 2002). Post-mortem examination of spleens and lymph nodes from patients with COVID-19 has revealed the lack of germinal centers, an essential part that can create a high-quality antibody response to produce long-term immunity (Kaneko et al., 2020). It is possible that TNF-α and IFN-γ signaling may aggravate lymphopenia through direct killing of lymphocytes. Indeed, severe COVID-19 cases have been reported to have massive amounts of pro-inflammatory cytokines including TNF-α in the germinal centers, thereby limiting the appropriate immune response.

Beyond the implications of our findings for patients with COVID-19, we also observed that TNF-α and IFN-γ–mediated pathology plays a critical role in other diseases and syndromes associated with cytokine storm. One key example is sepsis. According to the WHO, 11 million sepsis-related deaths occur worldwide each year (Rudd et al., 2020). Cytokine storm is a key component of sepsis pathology early in the course of disease, and several clinical trials have evaluated the efficacy of blocking individual cytokines without success (Chousterman et al., 2017), although a subsequent meta-analysis of trials using anti-TNF-α therapies has indicated some therapeutic benefit (Qiu et al., 2013). Our findings suggest that synergy between TNF-α and IFN-γ may be playing a key role in this pathogenesis. Similarly, hemophagocytic lymphohistiocytosis, which encompasses several related diseases including familial HLH, secondary HLH following infection, and autoimmune-associated macrophage activation syndrome (MAS), is characterized by a hyperactive immune response and cytokine storm and is often fatal (George, 2014). Emapalumab, an anti-IFN-γ antibody, was recently approved by the FDA to treat refractory, recurrent, or progressive HLH (Locatelli et al., 2020), and our data suggest that the therapeutic potential of combining this with an anti–TNF-α antibody could be beneficial. Additionally, cytokine storm has been reported in several other bacterial and viral infections including group A streptococcus, H5N1 virus, influenza H1N1 virus, SARS-CoV, and MERS-CoV (Channappanavar and Perlman, 2017; de Jong et al., 2006; Liu et al., 2016; Tisoncik et al., 2012). Elevated cytokine levels also likely play a role in the clinical pathology of multisystem inflammatory syndrome in children (MIS-C), characterized by shock, cardiac dysfunction, and gastrointestinal symptoms following infection with SARS-CoV-2 (Cheung et al., 2020; Jiang et al., 2020). Furthermore, numerous inflammatory syndromes, including manifestations of MAS, graft versus host disease, and the inflammation that results from cancer immunotherapies, also cause cytokine storm. Therefore, our studies defining the TNF-α and IFN-γ signaling pathway and the mechanistic dissection of this process and its induction of inflammatory cell death, PANoptosis, could potentially be applied in those conditions. Moreover, our study provides a mechanistic basis to define cytokine storm as a form of systemic inflammation initiated by cytokine-mediated inflammatory cell death, PANoptosis, which leads to the release of cellular contents including inflammatory cytokines and alarmins to propagate and exacerbate the systemic response.

Overall, identification of this critical inflammatory cell death pathway downstream of TNF-α and IFN-γ provides multiple druggable targets that can be examined for their efficacy in COVID-19 as well as other infectious and inflammatory diseases that involve cytokine storm and TNF-α and IFN-γ. With the failure of several high-profile clinical trials and the mounting COVID-19 morbidity and mortality, our discovery of the mechanisms involved in disease pathology during COVID-19 and identification of several therapeutic targets in the TNF-α and IFN-γ–mediated cell death pathway, PANoptosis, fills a critical unmet need and serves as a foundation for the development of evidence-based therapeutic strategies to mitigate this ongoing public health crisis.

### Limitations of Study and Future Directions

Our study tested the combination of anti-TNF-α and anti-IFN-γ neutralizing antibodies in SARS-CoV-2 infection, cytokine shock, sepsis, and HLH in mice. The degree of protection provided by this combination therapy in the SARS-CoV-2 mouse model was lower than that observed in the other non-infection cytokine shock models. We administered the treatment starting on day 1 after SARS-CoV-2 infection, but earlier administration of the therapy may provide better protection. Also, while monotherapy with these agents was significantly less effective than the combination in sepsis and HLH models, the ability of monotherapy to prevent pathology during COVID-19 cannot be ruled out. Furthermore, examining the levels of TNF-α and IFN-γ, as well as the inflammatory cell death, in the presence and absence of therapy in the SARS-CoV-2 mouse model will be important for the future.

In addition to TNF-α and IFN-γ, each molecule in the signaling pathway that we identified, including JAK and caspases, could potentially be targeted to prevent cytokine storm and should be investigated in the context of COVID-19. However, not all the molecules in the pathway have FDA-approved inhibitors ready for clinical usage. JAK inhibitors are approved for inflammatory diseases and have been shown to improve COVID-19 symptoms. Emricasan, an FDA-approved PAN-caspase inhibitor, has not been tested for COVID-19 but can be considered. Overall, to progress from fundamental discovery to therapeutic benefit, it will be important to extend the therapies identified in this study to humans. Adequately powered randomized controlled trials using the anti-TNF-α and anti-IFN-γ combination, JAK inhibitors, or PAN-caspase inhibitors administered at different timepoints in the disease course of COVID-19 are crucial to fully understand the efficacy of these treatments.

## ACKNOWLEDGEMENTS

We thank all the members of the Kanneganti laboratory for their comments and suggestions during the development of this manuscript. We thank A. Burton (St. Jude Children’s Research Hospital) for her technical support. We also thank R. Tweedell, PhD, for scientific editing and writing support. Work from our laboratory is supported by the US National Institutes of Health (AI101935, AI124346, AR056296, and CA253095 to T.-D.K.) and the American Lebanese Syrian Associated Charities (to T.-D.K.). The content is solely the responsibility of the authors and does not necessarily represent the official views of the National Institutes of Health. The *Irf3^-/-^, Irf7^-/-^*, and *Irf9^-/-^* mutant mice were provided by RIKEN BRC through the National Bio-Resource Project of the MEXT, Japan. We thank V.M. Dixit and N. Kayagaki (Genentech) for the *Casp11^-/-^* and *Casp1^-/-^ Casp11^-/-^* mutant mouse strains. We thank Dr. Michael Gale for the *Mavs^-/-^* mutant mouse strain. We thank K. Pfeffer for the *Gbp2^-/-^* mutant mouse strain. The following reagent was deposited by the Centers for Disease Control and Prevention and obtained through BEI Resources, NIAID, NIH: SARS-Related Coronavirus 2, Isolate USA-WA1/2020, NR-52281. The graphical abstract was created using BioRender.

## AUTHOR CONTRIBUTIONS

R.K. and T.-D.K. conceptualized the study; R.K. and B.R.S designed the methodology; R.K., B.R.S., S.T., E.P.W., L.Z., M.Z., B.S., B.B., and R.K.S.M. performed the experiments; P.Samir, P. Schreiner, and G.N. conducted the gene expression and publicly available dataset analysis; C.B.J. and R.W. directed the SARS-CoV-2 infections and provided scientific discussion; P.V. conducted the immunohistochemistry and pathology analysis; R.K., and T.-D.K. wrote the manuscript with input from all the authors. T.-D.K. acquired the funding and provided overall supervision.

## DECLARATION OF INTERESTS

St. Jude Children’s Research hospital filed a provisional patent application on TNF-α and IFN-γ signaling described in this study, listing R.K. and T.-D.K. as inventors (Serial No. 63/106,012).

## SUPPLEMENTAL FIGURE LEGENDS

**Figure S1. Co-treatment of TNF-α and IFN-γ induces cell death, Related to Figure 1**

(A) Percent of bone marrow-derived macrophages (BMDMs) that are dead 48 h after cytokine treatment using the IncuCyte imaging system and propidium iodide (PI) staining. BMDMs were stimulated with the indicated cytokines, “Cocktail-1” (IL-6, IL-18, IFN-γ, IL-15, TNF-α, IL-1α, IL-1β, and IL-2), “Cocktail-2” (IL-6, IL-18, IL-15, IL-1α, IL-1β, and IL-2), Cocktail-2+TNF-α, or Cocktail-2+IFN-γ. (B) Percent of BMDMs that are dead 48 h after treatment with the indicated cytokines using the IncuCyte imaging system and PI staining. (C) Percent of BMDMs that are dead 48 h after treatment with increasing concentration of cytokines in Cocktail-2 or TNF-α and IFN-γ using the IncuCyte imaging system and PI staining. (D) Real-time analysis of cell death in BMDMs using the IncuCyte imaging system and propidium iodide (PI) staining after treatment with increasing concentrations of TNF-α and IFN-γ. (E) Real-time analysis of cell death in primary human umbilical vein endothelial cells (HUVEC) treated with the indicated cytokines using the IncuCyte imaging system and PI staining. (F) Circulating amounts of TNF-α and IFN-γ in healthy volunteers and patients with mild, moderate, or severe COVID-19 (Silvin et al., 2020). (G) Expression of pro-inflammatory cytokines in macrophages, NK cells, CD8^+^ T cells, and B cells based on publicly available single-cell RNA-seq data using peripheral blood mononuclear cells obtained from healthy donors and patients with mild and severe COVID-19 (Lee et al., 2020b). Data are representative of at least three independent experiments. \*\**P* < 0.01; \*\*\*\**P* < 0.0001. Analysis was performed using the one-way ANOVA (A–C) or the two-way ANOVA (E and G). Data are shown as mean ± SEM.

**Figure S2. TNF-α and IFN-γ shock induces inflammatory responses and intestinal and lung damage, Related to Figure 2**

(A) CD45 immuno-staining in the intestine collected from mice injected intraperitoneally with PBS or TNF-α and IFN-γ at 5 h post-treatment. (B) Hematoxylin and eosin staining (H/E), cleaved caspase-3 (Clvd CASP3), and CD45 immuno-staining in the lungs collected from mice injected intraperitoneally with PBS or TNF-α and IFN-γ at 5 h post-treatment. Red arrows indicate stained cells for Clvd CASP3. (C) Quantitative analysis of Clvd CASP3-positive and TUNEL-positive cells in the intestine collected from mice injected intraperitoneally with PBS or TNF-α and IFN-γ at 5 h post-treatment. Fifty fields were analyzed under the microscope. (D) Quantitative analysis of Clvd CASP3-positive cells in the lungs collected from mice injected intraperitoneally with PBS or TNF-α and IFN-γ at 5 h post-treatment. Fifty fields were analyzed under the microscope. Data are representative of at least three independent experiments. Data are shown as mean ± SEM (C and D). \*\*\*\**P* < 0.0001. Analysis was performed using the *t* test (C and D).

**Figure S3. IRF1 and STAT1 are required for cell death downstream of TNF-α and IFN-γ cotreatment, Related to Figure 4**

(A) Percent of bone marrow-derived macrophages (BMDMs) that are dead 48 h after TNF-α and IFN-γ co-treatment using the IncuCyte imaging system and propidium iodide (PI) staining. (B) Real-time analysis of cell death in wild type (WT), *Irf1^-/-^*, and *Stat1^-/-^* BMDMs using the IncuCyte imaging system and PI staining after treatment with TNF-α and IFN-γ. (C) Representative images of cell death in WT, *Irf1^-/-^*, and *Stat1^-/-^* BMDMs are shown at 0 h and after 48 h of TNF-α and IFN-γ treatment. Scale bar, 50 μm. (D and E) Immunoblot analysis of (D) pro- (P35) and cleaved caspase-3 (P19 and P17; CASP3), pro- (P35) and cleaved caspase-7 (P20; CASP7), and pro- (P55) and cleaved caspase-8 (P18; CASP8) and (E) pro- (P53) and activated (P34) gasdermin E (GSDME), phosphorylated MLKL (pMLKL), and total MLKL (tMLKL) in WT and *Irf1^-/-^* BMDMs cotreated with TNF-α and IFN-γ for the indicated time. GAPDH was used as the internal control. Asterisks denote a nonspecific band. Data are representative of at least three independent experiments. \*\*\*\**P* < 0.0001. Analysis was performed using the one-way ANOVA (A) or two-way ANOVA (B). Data are shown as mean ± SEM (A and B).

**Figure S4. Nitric oxide produced downstream of IRF1 and STAT1 is required for cell death triggered by TNF-α and IFN-γ co-treatment, Related to Figure 4**

(A) Immunoblot analysis of iNOS in wild type (WT) and *Stat1^-/-^* bone marrow-derived macrophages (BMDMs) co-treated with TNF-α and IFN-γ for the indicated time. GAPDH was used as the internal control. (B) Nitric oxide production in WT, *Irf1^-/-^*, and *Stat1^-/-^* BMDMs co-treated with TNF-α and IFN-γ for the indicated time. (C) Immunoblot analysis of iNOS in WT and *Nos2^-/-^* BMDMs co-treated with TNF-α and IFN-γ for the indicated time. GAPDH was used as the internal control. (D) Production of nitric oxide by WT BMDMs treated with nitric oxide inhibitors, 1400W (100 μM) or L-NAME (1 mM), together with TNF-α and IFN-γ for 36 h. (E) Real-time analysis of cell death in PBS-, 1400W-, or L-NAME–treated WT BMDMs using the IncuCyte imaging system and propidium iodide (PI) staining after stimulation with TNF-α and IFN-γ. (F) Representative images of cell death in PBS-, 1400W-, or L-NAME–treated WT BMDMs are shown at 0 h and after 48 h of TNF-α and IFN-γ treatment. Scale bar, 50 μm. (G and H) Immunoblot analysis of (G) pro- (P35) and cleaved caspase-3 (P19 and P17; CASP3), pro- (P35) and cleaved caspase-7 (P20; CASP7), and pro- (P55) and cleaved caspase-8 (P18; CASP8) and (H) pro- (P53) and activated (P34) gasdermin E (GSDME), phosphorylated MLKL (pMLKL), and total MLKL (tMLKL) in WT and *Nos2^-/-^* BMDMs co-treated with TNF-α and IFN-γ for the indicated time. GAPDH was used as the internal control. Asterisk denotes a nonspecific band. Data are representative of at least three independent experiments. \*\*\*\**P* < 0.0001. Analysis was performed using the one-way ANOVA (D) or two-way ANOVA (B and E). Data are shown as mean ± SEM (B, D, and E).

**Figure S5. Concentration of nitric oxide is critical to induce cell death, and IFN-γ does not suppress TNF-α–mediated NF-κB signaling, Related to Figure 4**

(A) Immunoblot analysis of iNOS in wild type (WT) bone marrow-derived macrophages (BMDMs) treated with TNF-α alone, IFN-γ alone, or TNF-α and IFN-γ together for 24 h. GAPDH was used as the internal control. (B) Nitric oxide production in WT BMDMs treated with TNF-α alone, IFN-γ alone, or TNF-α and IFN-γ together for the indicated time. (C) Real-time analysis of cell death in PBS- and nitric oxide donor SIN-1–treated WT BMDMs using the IncuCyte imaging system and propidium iodide (PI) staining. (D and E) Heatmap depicting the expression levels of NF-κB target genes for (D) inflammatory cytokines/chemokines and (E) apoptosis regulators in WT BMDMs treated with TNF-α alone or co-treated with TNF-α and IFN-γ for 16 h relative to their expression in untreated (Mock) BMDMs. Data are representative of at least three independent experiments. \*\*\**P* < 0.001; \*\*\*\**P* < 0.0001. Analysis was performed using the two-way ANOVA (B and C).

**Figure S6. FADD regulates cell death induced by TNF-α and IFN-γ co-treatment, Related to Figure 5**

(A) Real-time analysis of cell death in wild type (WT), *Ripk3^-/-^* and *Ripk3^-/-^ Fadd^-/-^* bone marrow-derived macrophages (BMDMs) using the IncuCyte imaging system and propidium iodide (PI) staining after co-treatment with TNF-α and IFN-γ. (B) Representative images of cell death in WT, *Ripk3^-/-^*, and *Ripk3^-/-^ Fadd^-/-^* BMDMs are shown at 0 h and after 48 h of TNF-α and IFN-γ treatment. Scale bar, 50 μm. (C) Immunoblot analysis of pro- (P55) and cleaved caspase-8 (P18; CASP8), pro- (P35) and cleaved caspase-3 (P19 and P17; CASP3), and pro- (P35) and cleaved caspase-7 (P20; CASP7) in WT and *Ripk3^-/-^ Fadd^-/-^* BMDMs co-treated with TNF-α and IFN-γ for the indicated time. GAPDH was used as the internal control. (D) Immunoblot analysis of pro- (P53) and activated (P34) gasdermin E (GSDME), phosphorylated MLKL (pMLKL), and total MLKL (tMLKL) in WT and *Ripk3^-/-^ Fadd^-/-^* BMDMs co-treated with TNF-α and IFN-γ for the indicated time. GAPDH was used as the internal control. Asterisk denotes a nonspecific band. Data are representative of at least three independent experiments. \*\*\*\**P* < 0.0001. Analysis was performed using the two-way ANOVA (A). Data are shown as mean ± SEM (A).

**Figure S7. Deletion of individual cell death pathways is not sufficient to protect cells from death induced by TNF-α and IFN-γ, Related to Figure 5**

(A-C) Real-time analysis of cell death in wild type (WT), *Casp7^-/-^*, and *Casp3^-/-^* bone marrow-derived macrophages (BMDMs) (A); WT, *Gsdmd^-/-^, Gsdme^-/-^, Mlkl^-/-^*, and *Gsdmd^-/-^ Gsdme^-/-^* (*Gsdmd/e^-/-^*)*Mlkl^-/-^* BMDMs (B); or WT, *Casp1^-/-^, Casp11^-/-^*, and *Casp1/11^-/-^* BMDMs (C) using the IncuCyte imaging system and propidium iodide (PI) staining after co-treatment with TNF-α and IFN-γ. Representative images of cell death are shown at 0 h and after 48 h of TNF-α and IFN-γ treatment. Scale bar, 50 μm. \*\*\*\**P* < 0.0001. Analysis was performed using the two-way ANOVA. Data are representative of at least three independent experiments. Data are shown as mean ± SEM.

## METHODS

### Resource Availability

#### Lead Contact

Further information and requests for reagents may be directed to, and will be fulfilled by the lead contact Thirumala-Devi Kanneganti.

#### Materials Availability

All unique reagents generated in this study are available from the Lead Contact.

#### Data and Code Availability

The microarray data can be accessed in GEO (GSE160163). Publicly available datasets were obtained from Lucas et al. (Lucas et al., 2020), Silvin et al. (Silvin et al., 2020) (E-MTAB-9221), Hadjadj et al. (Hadjadj et al., 2020), and Lee et al. (Lee et al., 2020b) (GSE149689). All other datasets generated or analyzed during this study are included in the published article.

### Experimental Model and Subject Details

#### Mice

*Irf1^-/-^* (Matsuyama et al., 1993), *Stat1^-/-^* (Durbin et al., 1996), *Ripk3^-/-^* (Newton et al., 2004), *Ripk3^-/-^ Fadd^-/-^* (Dillon et al., 2012), *Ripk3^-/-^ Casp8^-/-^* (Oberst et al., 2011), *Apaf1^-/-^* (JAX, #004373) (Honarpour et al., 2000), *Nos2^-/-^* (JAX, #002609) (Laubach et al., 1995), *Casp7^-/-^* (Lakhani et al., 2006), *Casp3^-/-^* (Zheng et al., 2000), *Gsdmd^-/-^* (Karki et al., 2018), *Gsdme^-/-^* (Skarnes et al., 2011), *Mlkl^-/-^* (Murphy et al., 2013), *Casp1^-/-^* (Man et al., 2016), *Casp11^-/-^* (Kayagaki et al., 2011), *Casp1/11^-/-^* (Kayagaki et al., 2011), *Irf2^-/-^* (Kayagaki et al., 2019), *Irf3^-/-^* (Sato et al., 2000), *Irf7^-/-^* (Honda et al., 2005), *Irf3^-/-^ Irf7^-/-^* (Karki et al., 2018), *Ifnar1^-/-^* (Muller et al., 1994), *Ifnar2^-/-^* (Fenner et al., 2006), *Irf9^-/-^* (Kimura et al., 1996), *Trif^-/-^* (Yamamoto et al., 2003), *Mda5^-/-^* (Kato et al., 2006), *Mavs^-/-^* (Suthar et al., 2010), *cGas^-/-^* (Schoggins et al., 2014), *Sting*^gt/gt^ (Sauer et al., 2011), *Gbp2^-/-^* (Degrandi et al., 2013), *Ptpn6^-/-^* (Croker et al., 2008) mice have been previously described. *Irf5^tm1Ppr^*/J mice were purchased from The Jackson Laboratory (Stock Number 017311) and crossed with mice expressing Cre recombinase to generate *Irf5^-/-^* mice. *Gsdmd^-/-^ Gsdme^-/-^* (*Gsdmd/e^-/-^*)*Mlkl^-/-^* mice were generated at our facility by crossing *Gsdme^-/-^, Gsdmd^-/-^*, and *Mlkl^-/-^* mice. *Irf1^-/-^ Irf2^-/-^* mice were generated at our facility by crossing *Irf1^-/-^* with *Irf2^-/-^* mice. For SARS-CoV-2 infections, K18-ACE-2 transgenic mice were purchased from The Jackson Laboratory (Stock Number 034860). All mice were generated on or extensively backcrossed to the C57/BL6 background.

All mice were bred at the Animal Resources Center at St. Jude Children’s Research Hospital and maintained under specific pathogen-free conditions. Both male and female mice were used in this study; age- and sex-matched 6-to 9-week old mice were used for *in vitro* and 7-to 8-week old mice were used for *in vivo* studies. Mice were maintained with a 12 h light/dark cycle and were fed standard chow. Non-infectious animal studies were conducted under protocols approved by the St. Jude Children’s Research Hospital committee on the Use and Care of Animals. SARS-CoV-2 infections were performed at the University of Tennessee Health Science Center under ABSL3 conditions in accordance with the approval of the Institutional Animal Care and Use Committee of University of Tennessee Health Science Center (Protocol #20-0132).

#### Cell culture

Primary mouse bone marrow-derived macrophages (BMDMs) were generated from the bone marrow of wild type and the indicated mutant mice. Cells were grown for 5–6 days in IMDM (Thermo Fisher Scientific, 12440053) supplemented with 1% non-essential amino acids (Thermo Fisher Scientific, 11140-050), 10% FBS (Biowest, S1620), 30% L929 conditioned media, and 1% penicillin and streptomycin (Thermo Fisher Scientific, 15070-063). BMDMs were then seeded into antibiotic-free media at a concentration of 1 x 10^6^ cells into 12-well plates and incubated overnight. The human monocytic cell line THP-1 (ATCC, TIB-202) was cultured in RPMI media (Corning, 10-040-CV) supplemented with 10% FBS and 1% penicillin and streptomycin. The primary umbilical vein endothelial cells from normal human (HUVEC) (ATCC, PCS-100-013) were cultured in vascular cell basal medium (ATCC, PCS-100-030) containing cell growth factors (ATCC, PCS-100-040) and 1% penicillin and streptomycin.

#### Isolation of PBMCs

Blood was collected from anonymous healthy donors at St. Jude Children’s Research Hospital following IRB-approved protocols. Human PBMCs were isolated from the blood by density gradient using Percoll (GE Healthcare, 17-5445-01). PBMCs were washed and resuspended in RPMI 1640 supplemented with 10% FBS.

#### SARS-CoV-2 culture

The SARS-CoV-2 isolate USA-WA1/2020 was obtained through BEI Resources (NIAID, NIH: SARS-Related Coronavirus 2, Isolate USA-WA1/2020, NR-52281) and amplified in Vero-E6 cells (ATCC, VERO C1008) at an MOI of 0.1 in Minimal Essential Medium (MEM; Corning, 17-305-CV) supplemented with 5% heat-inactivated FBS (Gibco) and 1% L-Glutamine (Corning, 25-005-Cl) and 5 mM penicillin/streptomycin (Gibco, 30-001-Cl). Following virus amplifications, viral titer was determined using a plaque assay using the method described previously for alphaviruses (Lee et al., 2020a). All experiments involving SARS-CoV-2 were done in a biosafety level 3 laboratory.

### Methods Details

#### Cell stimulation

BMDMs were stimulated with the following cytokines where indicated unless otherwise noted: 20 ng/mL of IL-6 (Peprotech, 212-16), 10 ng/mL of IL-18 (BioLegend, 767004), 20 ng/mL of IL-15 (R&D, 447-ML), 20 ng/mL of IL-1α (Peprotech, 200-01A), 20 ng/mL of IL-1β (R&D, 201-LB-025), 20 ng/mL of IL-2 (Peprotech, 212-12), 25 ng/mL of TNF-α (Peprotech, 315-01A), 50 ng/mL of IFN-γ (Peprotech, 315-05), 50 ng/mL of IFN-α (PBL Assay, 12106-1), 50 ng/mL of IFN-β (PBL Assay, 12400-1), or 50 ng/mL of IFN-λ (Peprotech, 250-33). For human cells, 50 ng/mL of TNF-α (Peprotech, AF-300-01A) and 100 ng/mL of IFN-γ (Peprotech, 300-02) was used for the indicated time. For the inhibition of NO, cells were co-treated with 1 mM of L-NAME hydrochloride (TOCRIS, 0665) or 100 μM of 1400W dihydrochloride (Enzo Life Sciences, ALX-270-073-M005). SIN-1 chloride (TOCRIS, 0750) at the indicated concentrations was used as the NO donor.

#### Real-time imaging for cell death

The kinetics of cell death were determined using the IncuCyte S3 (Essen BioScience) live-cell automated system. BMDMs (5 × 10^5^ cells/well) were seeded in 24-well tissue culture plates. THP-1 (1 × 10^5^ cells/well) and HUVEC (5 × 10^4^ cells/well) were seeded in 48-well tissue culture plates. Cells were treated with the indicated cytokines and stained with propidium iodide (PI; Life Technologies, P3566) following the manufacturer’s protocol. The plate was scanned, and fluorescent and phase-contrast images (4 image fields/well) were acquired in real-time every 1 h from 0 to 48 h post-treatment. PI-positive dead cells are marked with a red mask for visualization. The image analysis, masking, and quantification of dead cells were done using the software package supplied with the IncuCyte imager.

#### Immunoblot analysis

Cell lysates and culture supernatants were combined in caspase lysis buffer (containing 1× protease inhibitors, 1× phosphatase inhibitors, 10% NP-40, and 25 mM DTT) and 4× sample loading buffer (containing SDS and 2-mercaptoethanol) for immunoblot analysis of caspases. For immunoblot analysis of signaling components, supernatants were removed, and cells were washed once with DPBS, followed by lysis in RIPA buffer and sample loading buffer. Proteins were separated by electrophoresis through 8–12% polyacrylamide gels. Following electrophoretic transfer of proteins onto PVDF membranes (Millipore, IPVH00010), nonspecific binding was blocked by incubation with 5% skim milk, then membranes were incubated with primary antibodies against: caspase-3 (Cell Signaling Technology [CST], #9662, 1:1000), cleaved caspase-3 (CST, #9661, 1:1000), caspase-7 (CST, #9492, 1:1000), cleaved caspase-7 (CST, #9491, 1:1000), caspase-8 (AdipoGen, AG-20T-0138-C100, 1:1000), cleaved caspase-8 CST, #8592, 1:1000), caspase-9 (CST, #9504, 1:1000), caspase-11 (Novus Biologicals, NB120-10454, 1:1000), caspase-1 (AdipoGen, AG-20B-0042, 1:1000), GAPDH (CST, #5174, 1:1000), iNOS (CST, #13120, 1:1000), pMLKL CST, #37333, 1:1000), tMLKL Abgent, AP14272b, 1:1000), pRIPK1 (CST, #31122, 1:1000), tRIPK1 (CST, #3493, 1:1000), GSDMD Abcam, ab209845, 1:1000), GSDME (Abcam, #19859, 1:1000), β-actin (Proteintech, 66009-1-IG, 1:1000). Membranes were then washed and incubated with the appropriate horseradish peroxidase (HRP)–conjugated secondary antibodies (Jackson ImmunoResearch Laboratories, anti-rabbit [111-035-047] 1:5000, anti-mouse [315-035-047] 1:5000, and anti-rat [112-035-003], 1:5000). Proteins were visualized using Immobilon Forte Western HRP Substrate (Millipore, WBLUF0500).

#### Nitric oxide measurement

The amount of nitric oxide released in the supernatant of BMDMs stimulated with TNF-α alone, IFN-γ alone, or TNF-α + IFN-γ was determined using Griess reagent kit (Invitrogen, G7921) according to the manufacturer’s instructions.

#### In vivo TNF-α and IFN-γ–induced inflammatory shock

Age- and gender-matched 6-to 8-week-old mice of the indicated genotypes were injected intraperitoneally with 10 μg TNF-α alone, 20 μg IFN-γ alone, or TNF-α + IFN-γ diluted in DPBS. Animals were under permanent observation, and survival was assessed every 30 min. Blood was collected 5 h after cytokine injection. Blood composition was analyzed using an automated hematology analyzer. Serum LDH, ALT, AST, blood urea nitrogen (BUN), and ferritin were analyzed by colorimetry using respective kits (LDH, REF#A11A01824; ALT, REF#A11A01627; AST, REF#A11A01629; and BUN, REF#A11A01641, all from HORIBA; and ferritin, ab157713, Abcam) according to the manufacturer’s instructions.

For neutralizing antibody treatment, age- and gender-matched 6-to 8-week-old WT mice were administered intraperitoneally 200 μl of DPBS containing 500 μg of isotype control (Leinco Technologies, Inc., I-536) (n = 10) or 500 μg of neutralizing antibody against TNF-α (Leinco Technologies, Inc., T-703) plus 500 μg of neutralizing antibody against IFN-γ (Leinco Technologies, Inc., I-1190) (n = 14) 12 h before i.p. injection of TNF-α and IFN-γ. Mice were monitored for survival. The results were pooled from 2 independent experiments.

#### Flow cytometry

Peripheral blood was mixed with ACK lysis buffer for 3 min to lyse the RBCs. The cells were then stained with the following monoclonal antibodies for flow cytometry: CD11b (M1/70; 48-0112-82) from eBioscience; F4/80 (BM8; 123116), Ly6C (HK1.4; 128016), and Ly6G (1A8; 127616) from BioLegend; and CD19 (1D3; 35-0193-U025), TCRb (H57-597; 60-5961-U100), and CD45.2 (104; 50-0454-U100) from Tonbo Biosciences. Cells were gated on live single-cell populations and hematopoietic cells using the CD45.2 gate followed by separation of each of the specific cell populations using the following cell surface markers: macrophages (CD11b^+^, F4/80^+^), neutrophils (CD11b^+^, Ly6C^low^, Ly6G^high^), T cells (TCRb^+^, CD19^-^), and B cells (CD19^+^, TCRb^-^).

#### Histopathology

Lungs and intestines were fixed in 10% formalin, then processed and embedded in paraffin by standard procedures. Sections (5 μM) were stained with hematoxylin and eosin (H&E) and examined by a pathologist blinded to the experimental groups. For immunohistochemistry, formalin-fixed paraffin-embedded lungs and intestines were cut into 4 μM sections. Cleaved caspase-3 (Essen Bioscience, 4704) and CD45 (BD Pharmingen™, 553076) staining was performed according to the manufacturer’s instructions. TUNEL (terminal deoxynucleotidyl transferase deoxyuridine triphosphate nick-end labeling) staining was performed using the DeadEnd kit (Promega, PRG7130) according to the manufacturer’s instructions. The number of cleaved caspase-3– and TUNEL-positive cells in five high power fields (20x) were counted per mouse.

#### COVID-19 patient cytokine analysis

COVID-19 patient cytokine data was obtained from Lucas et al. (Lucas et al., 2020). The list of the top 10 differentially present pro-inflammatory cytokines was manually generated. Amounts of selected cytokines were averaged for healthy control subjects, and moderate and severe COVID-19 patients. Heatmaps were generated using Morpheus (https://software.broadinstitute.org/morpheus). Cytokine data for the serum levels of TNF-α and IFN-γ was kindly shared by Silvin et al. (Silvin et al., 2020) (E-MTAB-9221) for healthy patients and patients with moderate, severe, and critical COVID-19.

#### Gene expression analysis

Genes involved in type II interferon signaling were selected based on annotations (Hadjadj et al., 2020). Nanostring nCounter data was kindly shared by Hadjadj et al. for healthy patients and patients with moderate, severe, and critical COVID-19. Heatmaps were generated based on average expression of selected genes using Morpheus. *NOS2* expression in COVID-19 patients was extracted from the same Nanostring nCounter data (Hadjadj et al., 2020).

Murine orthologs of the genes identified as the most differentially regulated in humans were manually selected from microarray-based transcriptomics analysis of BMDMs treated with TNF-α and IFN-γ and used to generate heatmaps using Morpheus. Additionally, NF-κB target genes for both inflammatory cytokines/chemokines and apoptosis regulators (https://www.bu.edu/nf-kb/gene-resources/target-genes/) were selected from the microarray of BMDMs.

#### Single-cell RNA-seq data analysis

The feature-barcode matrix was downloaded from GSE149689 (Lee et al., 2020b) (GSE149689). Low quality cells were excluded from the analysis if the mitochondrial genes represented > 15% or if the number of features in a cell was < 200. Count values were normalized, log-transformed, and scaled using the Seurat R package v3.1.4 (Satija et al., 2015). The top 15 dimensions and a resolution value of 0.5 were used for UMAP dimension reduction (Lee et al., 2020b).

#### In vitro infection of PBMCs

For SARS-CoV-2 infection, 1 × 10^6^ PBMCs from 3 healthy donors were seeded into 12-well plates and infected with indicated MOI of the virus. Supernatants from medium-treated or infected PBMCs were collected at 24 h post-infection. The pro-inflammatory cytokines in the supernatant were analyzed by multiplex ELISA (Millipore, HCYTMAG-60K-PX29) according to the manufacturer’s instructions.

#### SARS-CoV-2 infection in mice

Age- and gender-matched, 7-to 8-week old K18-ACE-2 transgenic mice were anesthetized with 5% isoflurane and then infected intranasally with SARS-CoV-2 in 50 μl DPBS containing around 2 × 10^4^ PFU. Infected mice were administered intraperitoneally 200 μl of DPBS containing 500 μg of isotype control (Leinco Technologies, Inc., I-536) (n = 12) or 500 μg of neutralizing antibody against TNF-α (Leinco Technologies, Inc., T-703) plus 500 μg of neutralizing antibody against IFN-γ (Leinco Technologies, Inc., I-1190) (n = 13) on days 1, 3, and 4 post-infection. Mice were monitored over a period of 14 days for survival.

#### LPS-induced sepsis

Age- and gender-matched, 7-to 8-week old WT mice were injected intraperitoneally with 20 mg/kg body wight of LPS (Sigma, L2630). The mice were administered intraperitoneally 200 μl of DPBS containing no antibody (n = 17), 500 μg of isotype control (n = 10), 500 μg of neutralizing antibody against TNF-α (n = 17), 500 μg of neutralizing antibody against IFN-γ (n = 17), or 500 μg each of neutralizing antibodies against TNF-α and IFN-γ (n = 18) 30 min and 6 h post-LPS injection. Mice were monitored over a period of 3 days for survival. The results were pooled from 2 independent experiments.

#### Poly I:C and LPS–induced hemophagocytic lymphohistiocytosis (HLH)

HLH was induced by sequential challenge of poly I: C and LPS as described previously (Wang et al., 2019). Age- and gender-matched, 7-to 8-week old WT mice were injected intraperitoneally with 10 mg/kg body wight of high molecular weight poly I:C (InvivoGen, tlrl-pic). The mice were then administered intraperitoneally 200 μl of DPBS containing no antibody (n = 15), 500 μg of isotype control (n = 15), 500 μg of neutralizing antibody against TNF-α (n = 15), 500 μg of neutralizing antibody against IFN-γ (n = 15), or 500 μg each of neutralizing antibodies against TNF-α and IFN-γ (n = 15) 24 h post-poly I:C injection. All the mice were then challenged with a sub-lethal dose of LPS (5 mg/kg body weight) 1 h after the above-mentioned treatments. Mice were monitored over a period of up to 3 days for survival. The results were pooled from 2 independent experiments.

### Quantification and Statistical Analysis

GraphPad Prism 8.0 software was used for data analysis. Data are shown as mean ± SEM. Statistical significance was determined by *t* tests (two-tailed) for two groups or one-way ANOVA or two-way ANOVA for three or more groups. Survival curves were compared using the log-rank (Mantel-Cox) test. *P* < 0.05 was considered statistically significant.

## Notes

### Competing Interest Statement

St. Jude Childrens Research hospital filed a provisional patent application on TNF-α and IFN-γ signaling described in this study, listing R.K. and T.-D.K. as inventors (Serial No. 63/106,012).

### Summary of Updates

Discussion updated to add limitations section.

