## Supplementary Figures 1-7 for "Synergism of TNF-α and IFN-γ triggers inflammatory cell death, tissue damage, and mortality in SARS-CoV-2 infection and cytokine shock syndromes"

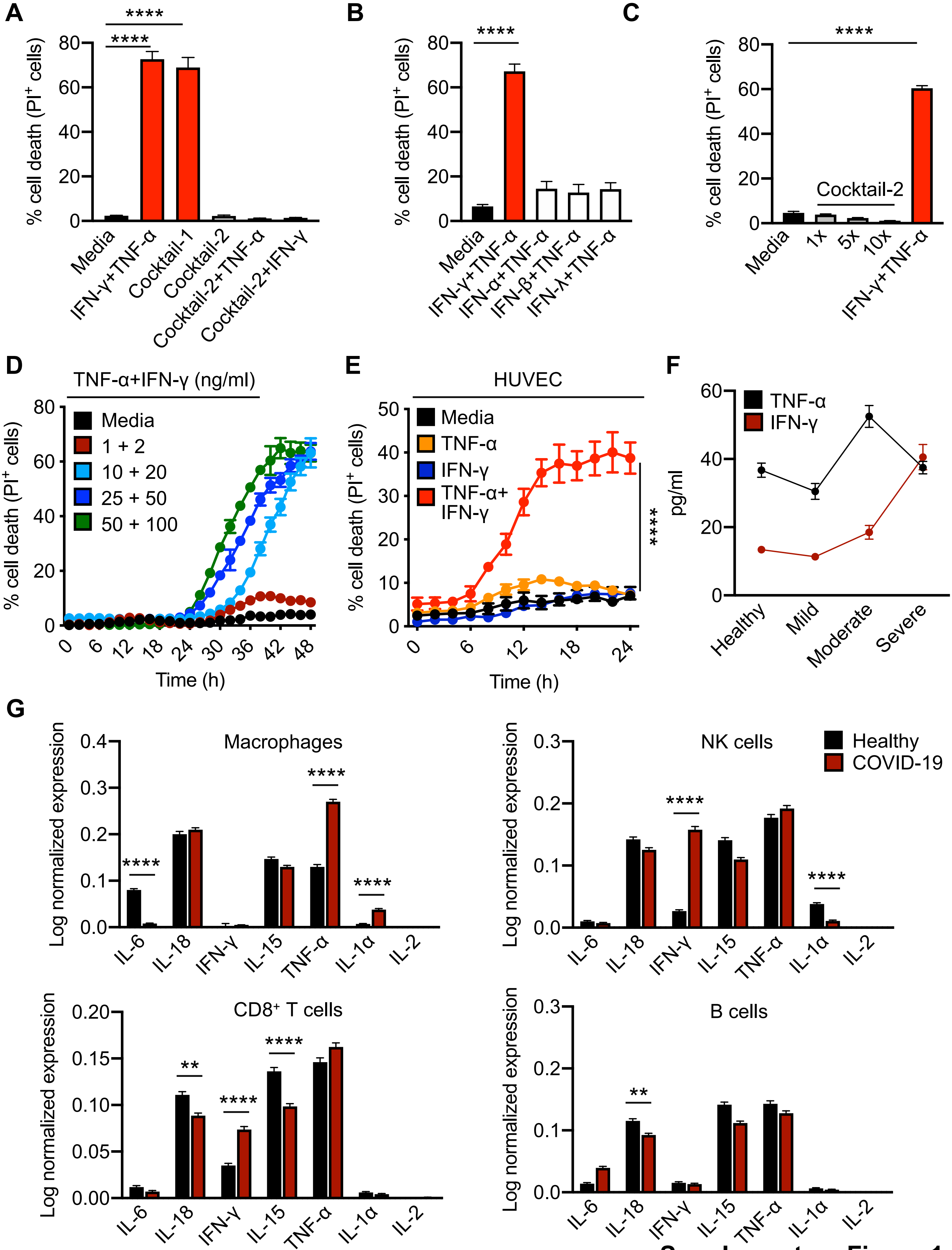


**Supplementary Figure 1**

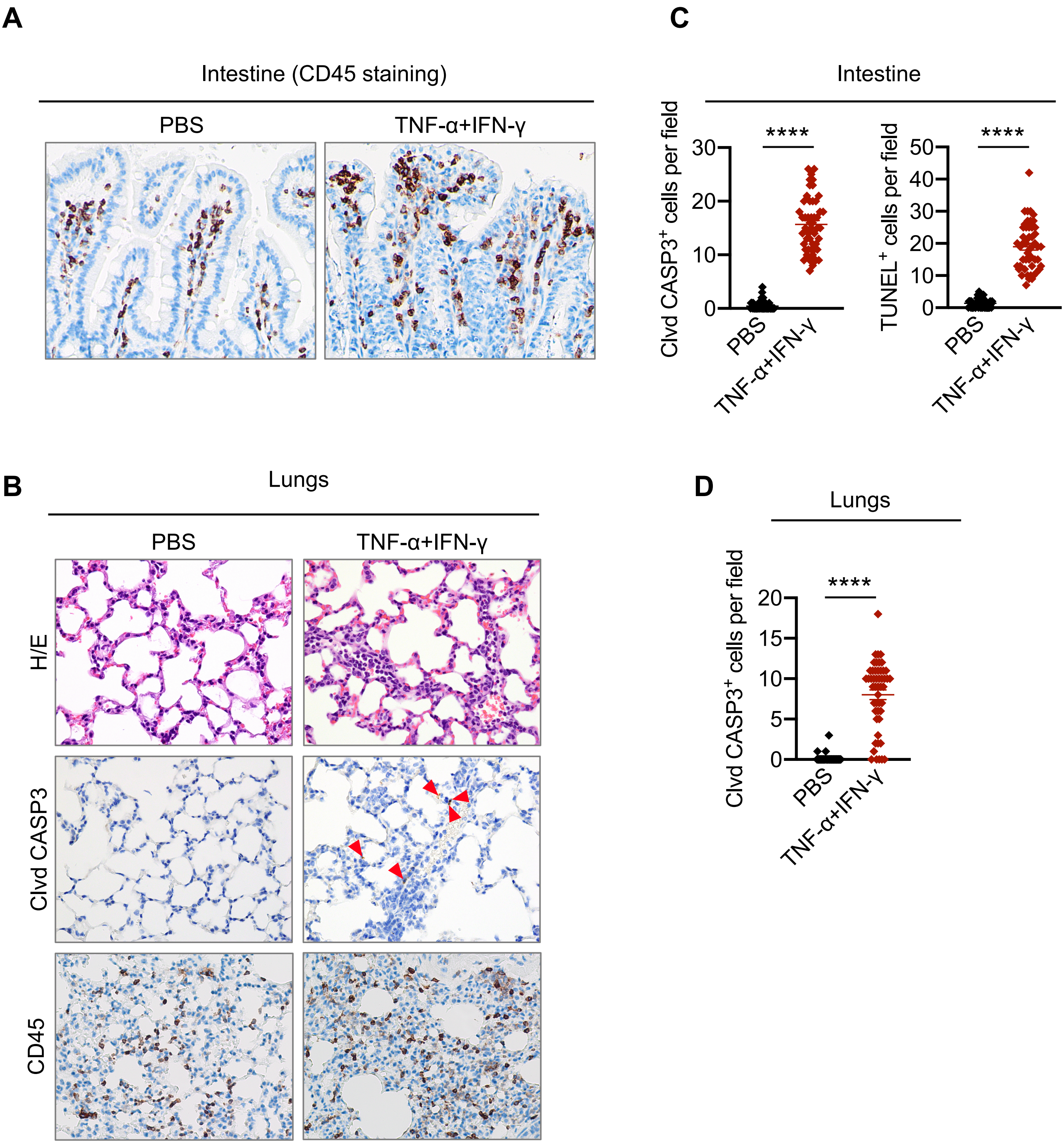


**Supplementary Figure 2**

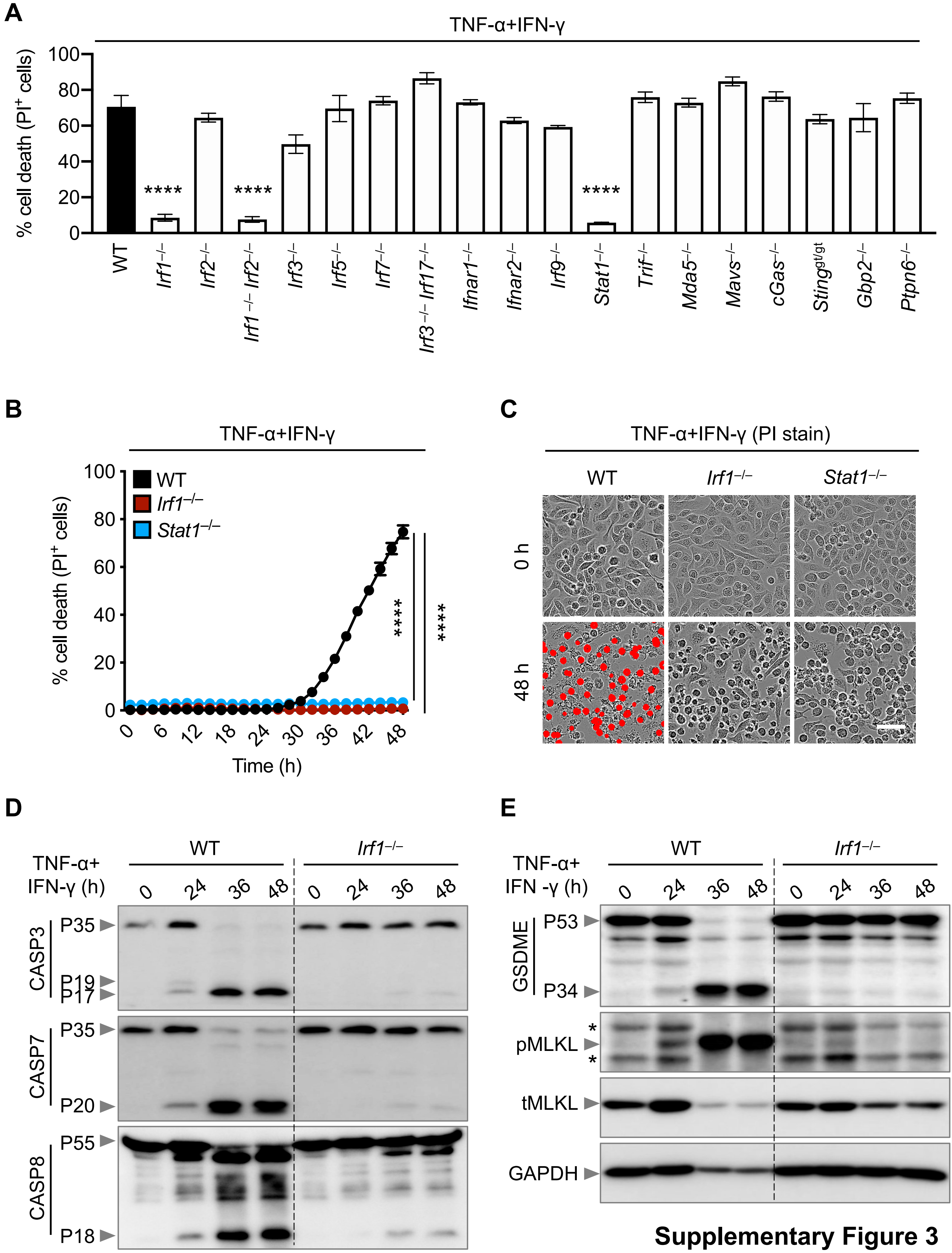


**Supplementary Figure 3**

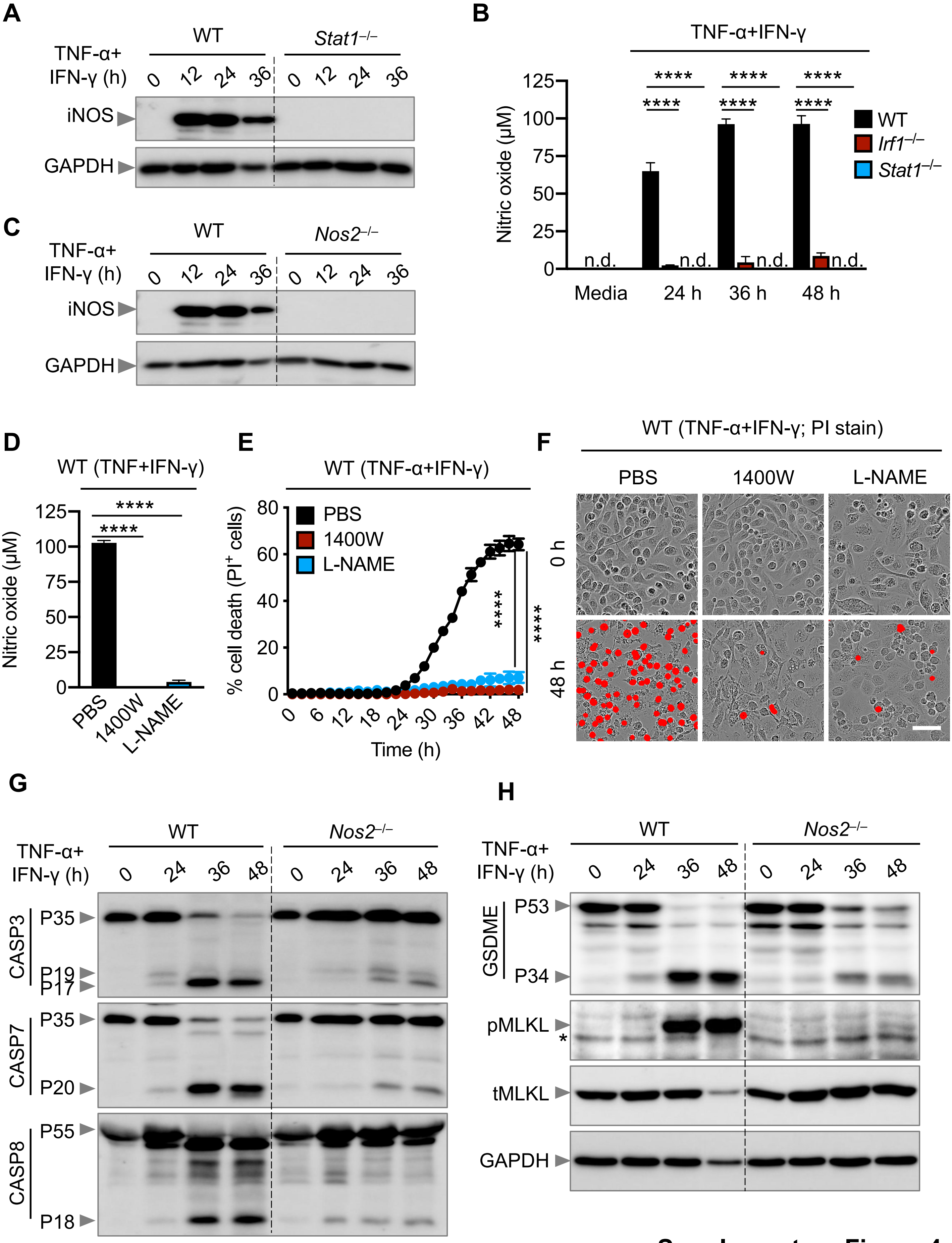


Supplementary Figure 4

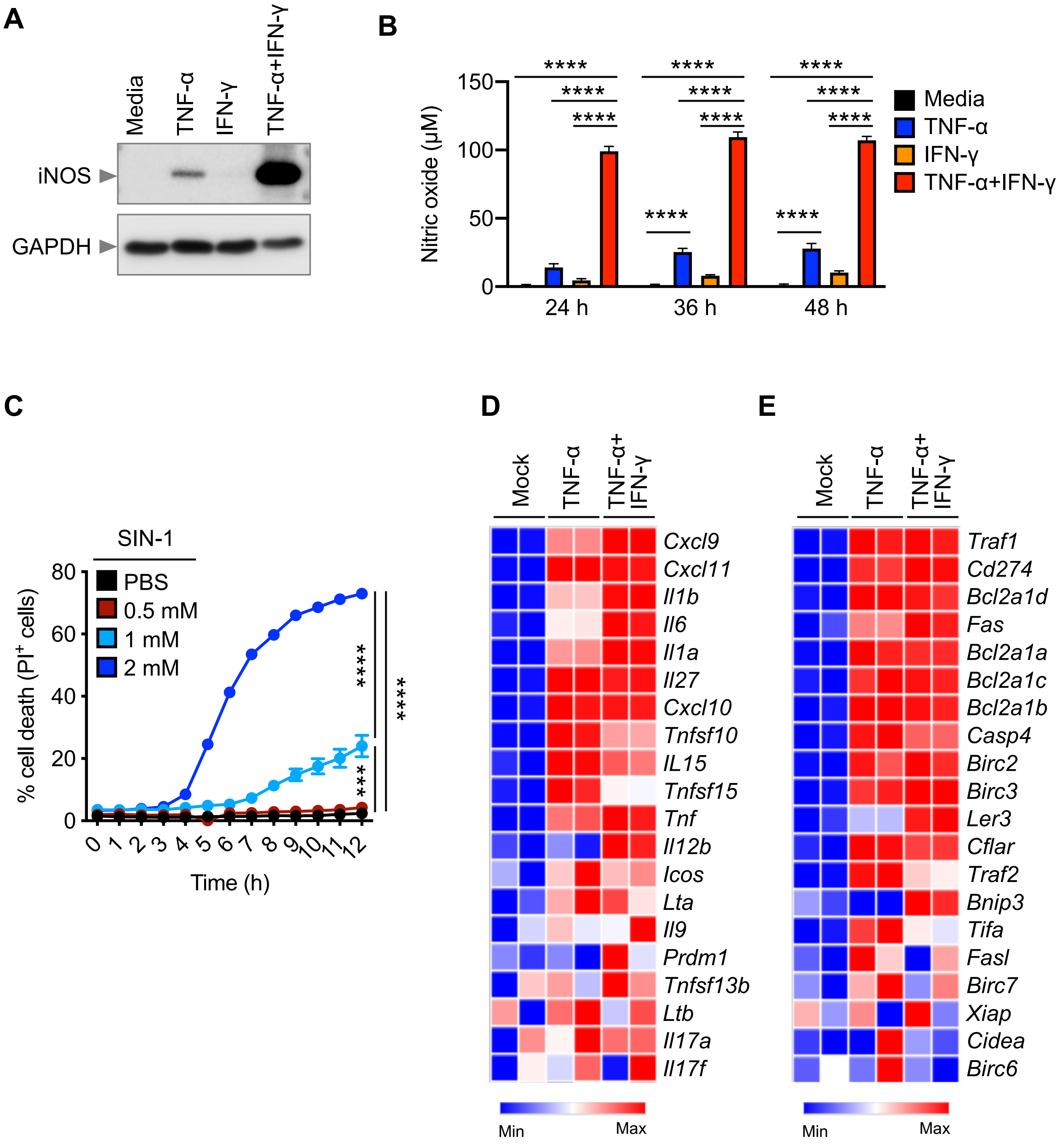


**Supplementary Figure 5**

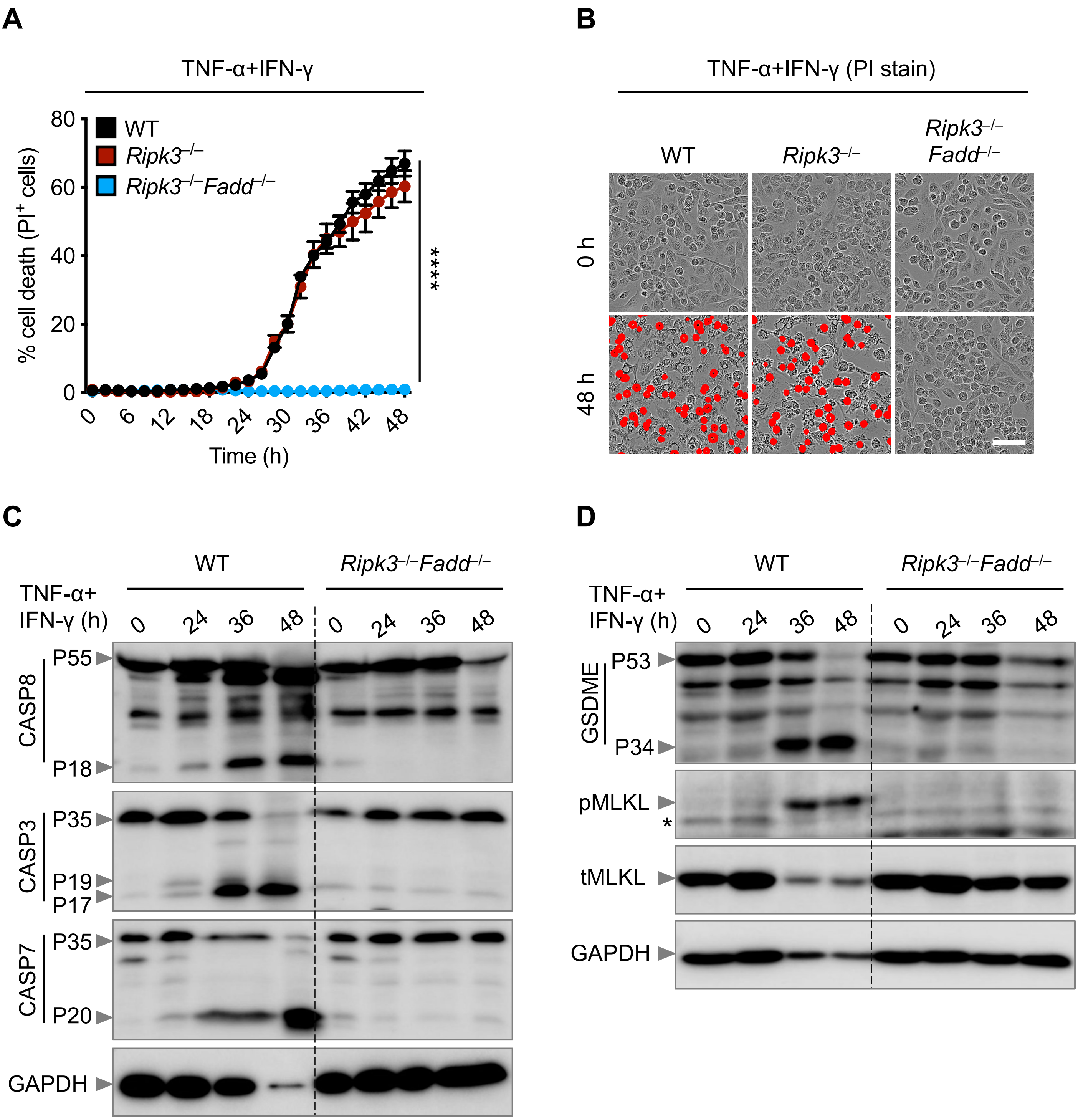


**Supplementary Figure 6**

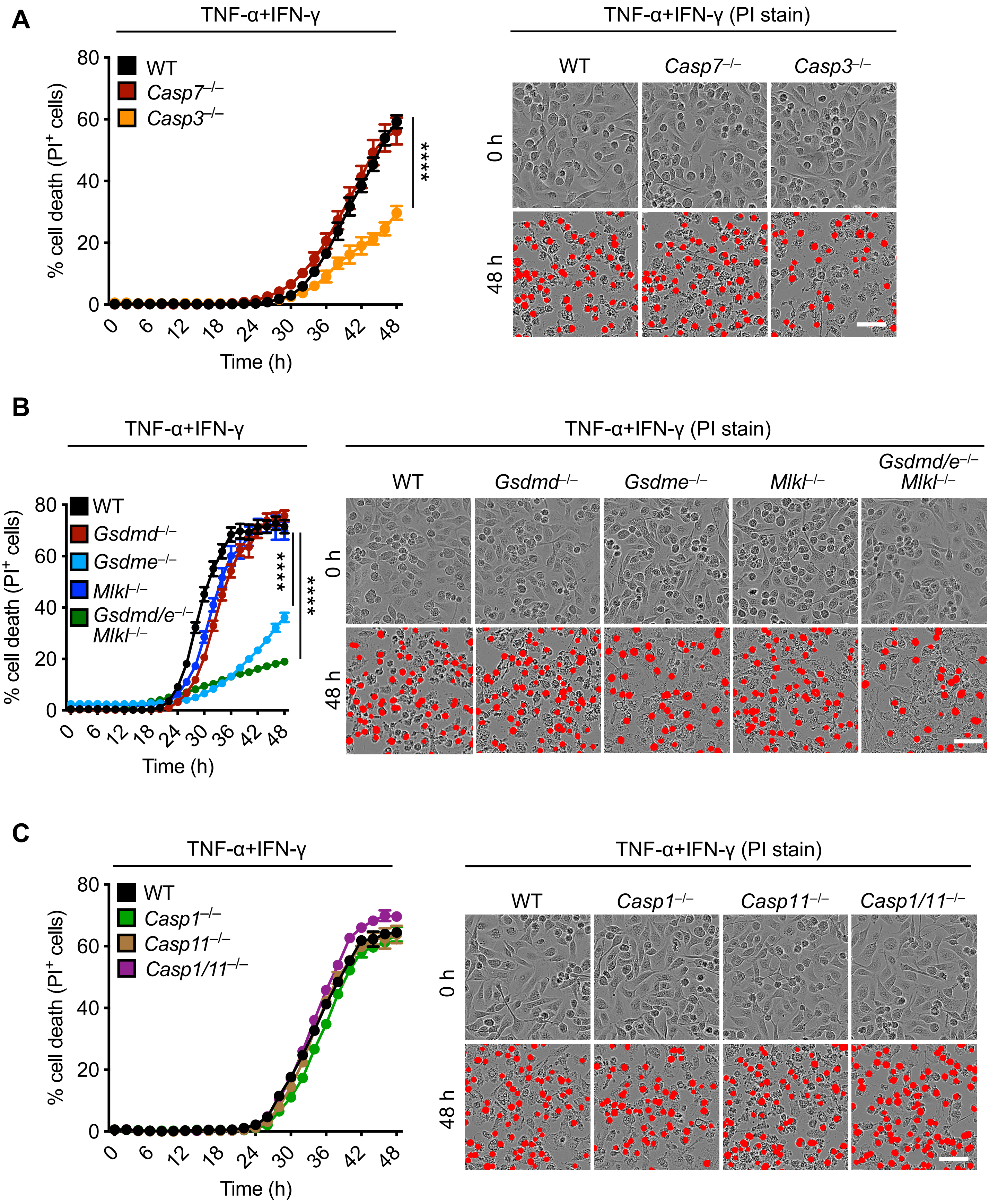


**Supplementary Figure 7**
